# Higher T-cell density in primary prostate cancer is associated with reduced fraction of CD8 effector cells and increased TIGIT

**DOI:** 10.64898/2026.08.27.747524

**Authors:** Seifeldin Awad, Carla Calagua, Olga S. Voznesensky, Sara Abdelkader, Razan Mohanna, Haydn Kissick, Sabina Signoretti, David J. Einstein, Steven P. Balk

## Abstract

A subset of untreated primary prostate cancer (PCa) contain substantial focal T-cell infiltrates, but whether these reflect antitumor responses that could potentially be enhanced by immune checkpoint blockade (ICB) remains unclear. We used immunohistochemistry, immunofluorescence, whole-slide spatial analysis, bulk RNA sequencing, and immune-cell deconvolution to characterize immune infiltrates in untreated primary PCa. Absolute CD8 T-cell density generally increased with total CD3 T-cell density, but the CD8/CD3 ratio decreased as overall T-cell density increased, indicating a preferential increase in CD4 T cells. Highly infiltrated tumors also had lower GZMB abundance relative to CD8 T-cell abundance. Multiplex analysis showed trends toward greater TIM3 and LAG3 expression among PD1⁺CD8⁺ T cells and increased regulatory T-cell features in highly infiltrated tumors. TIGIT⁺ cell density and the TIGIT/CD3 ratio increased with T-cell infiltration, whereas PD1/CD3 was not associated with overall CD3⁺ T-cell density. Both TIGIT/CD3 and PD1/CD3 ratios were enriched within lymphoid aggregates compared with matched tumor and benign regions, consistent with these structures being checkpoint-rich immune niches. Transcriptomic analyses supported a shift in relative immune composition toward CD4 T cells and selective increases in immune checkpoints. Together these findings suggest that effective immune responses in a subset of primary PCa with increased T-cell infiltration are being repressed by several mechanisms and may respond to therapies targeting specific immunosuppressive mechanisms.

## Introduction

Prostate cancer (PCa) is one of the most commonly diagnosed malignancies in men and a major cause of cancer-related mortality [1]. Immune checkpoint blockade (ICB) has demonstrated activity in molecularly mismatch-repair-deficient PCa [2], but has limited efficacy in unselected patients [3]. This limited activity is associated with a relatively immune-cold tumor microenvironment with sparse immune-cell infiltration, which may reflect low tumor immunogenicity and multiple immunosuppressive mechanisms [4,5]. Nevertheless, analyses of untreated radical prostatectomy specimens have demonstrated substantial heterogeneity in the abundance, composition, and spatial distribution of immune cells, indicating that a subset of primary tumors may be stimulating immune responses [6].

Detailed tissue studies have shown that immune-cell infiltration in primary PCa does not necessarily correspond to effective cytotoxic immunity. Multiplex immunohistochemical and whole-slide analyses have demonstrated that CD4⁺ T cells are frequently more abundant than CD8⁺ T cells and that both populations are concentrated predominantly in stromal rather than epithelial compartments. Cytotoxic T cells may remain relatively depleted from the tumor epithelium even when regulatory T cells, B cells, and macrophages accumulate in cancer-associated tissue. Many infiltrating T cells express PD1, and their abundance and localization vary according to tumor characteristics, including PTEN and ERG status [8,9]. Increased regulatory T-cell and M2-like macrophage infiltration has also been associated with biochemical recurrence.

Other studies have similarly demonstrated that immune infiltration in primary PCa is compositionally diverse and not uniformly associated with favorable outcomes. Macrophages can constitute a dominant immune population, and higher-grade tumors may contain greater densities of both CD8⁺ and CD4⁺ T cells without a corresponding association with biochemical recurrence. Although PD-L1⁺ cell densities are generally low, greater peritumoral PD-L1⁺ cell density has been independently associated with recurrence [10]. High-dimensional imaging has further identified recurrent cellular neighborhoods composed of malignant, immune, and stromal populations, including spatial associations among mast cells, M2-like macrophages, and regulatory T cells. These neighborhoods are influenced by androgen receptor-positive stromal cells and inflammatory gene-expression networks [11]. Collectively, these observations indicate that the biological significance of T-cell infiltration depends on its cellular, stromal, and myeloid context and cannot be inferred from immune-cell density alone.

The spatial organization of immune cells may provide additional information. Two T-cell patterns have been described in primary PCa: a clustered immune profile characterized by dense CD4⁺ T-cell aggregates interacting with PD-L1⁺ cells and a free immune profile characterized by CD8⁺ T cells positioned near tumor cells. The free immune profile was enriched in tumors with germline homologous-recombination repair defects and was associated with favorable pathological and clinical features [12]. Integrated single-cell RNA sequencing and spatial transcriptomics have further identified suppressive myeloid populations, exhausted T-cell states, regulatory T-cell interactions, and angiogenic stromal programs in untreated localized tumors, supporting the presence of coordinated immunosuppressive networks [13]. Tertiary lymphoid structures and less-organized lymphoid aggregates have also been identified in radical prostatectomy specimens and associated with increased T-cell, B-cell, major histocompatibility complex, and antigen-presentation gene-expression signatures [14,15]. However, the functional significance of these structures and their relationship to immune-checkpoint expression remain incompletely defined.

We previously identified and characterized a subset of untreated localized prostate cancers containing focal regions with substantial immune-cell infiltration [6]. Multiplex immunofluorescence demonstrated increased densities of CD8⁺ T cells within these foci, including substantial fractions expressing TIM3 and/or LAG3. We also identified cells that were negative for both markers and positive for TCF1, consistent with a stem-like or progenitor population that may retain responsiveness to ICB [7]. These findings suggested that immune-infiltrated tumors contain antigen-experienced T-cell populations at different states of differentiation. However, whether tumors with focal immune infiltration constitute a biologically distinct subset and whether their infiltrates reflect effective antitumor immunity or an increasingly checkpoint-regulated immune state remained unclear.

To address these questions, we integrated quantitative and spatial tissue analyses with bulk RNA sequencing and immune-cell deconvolution. We examined how increasing T-cell density relates to CD4/CD8 composition, granzyme B (GZMB)-associated cytotoxic differentiation, regulatory T-cell features, checkpoint expression, immune-cell composition, and the spatial organization of lymphoid aggregates. Although highly infiltrated tumors contained greater absolute numbers of CD8⁺ T cells, the proportional increase in non-CD8⁺ T cells was greater, resulting in reduced CD8⁺ T-cell dominance. Increasing infiltration was also associated with lower GZMB abundance relative to CD8⁺ T-cell abundance and increased TIGIT expression. Transcriptomic analyses further supported altered immune composition and broader immune-regulatory and checkpoint-associated programs. Together, these findings indicate that focal T-cell infiltration in untreated primary PCa represents an endogenous immune response accompanied by altered immune composition and checkpoint adaptation.

## Methods

### Patient cohort and tissue specimens

This study included 54 archival formalin-fixed, paraffin-embedded specimens of untreated primary PCa. Cases were selected based on the availability of adequate tumor-containing tissue for histopathological assessment, chromogenic immunohistochemistry, multiplex immunofluorescence, and quantitative digital analysis. The cohort encompassed a broad range of immune infiltration, permitting comparison of immune-rich and immune-poor tumors. Gleason scores were available for 52 specimens and ranged from 6 to 10, with most specimens having a Gleason score of 7 (n = 41). Specimens were processed according to routine clinical pathology procedures, and serial tissue sections were used for hematoxylin and eosin staining, immunohistochemistry, and multiplex immunofluorescence. Whole-tumor tissue and, where available, histologically benign prostate tissue, immune-infiltrated tumor regions, and lymphoid aggregates were evaluated separately. Matched tumor and benign regions were available for 25 specimens. The number of evaluable specimens varied among assays because of differences in tissue availability, staining quality, and availability of the required measurements. In accordance with institutional policy governing the use of these archival specimens, institutional review board approval and individual informed consent were not required.

### Histopathological assessment

Tissue sections were reviewed by a pathologist (Seifeldin Awad) to identify prostate tumor, histologically benign prostate tissue, immune-infiltrated tumor foci, and lymphoid aggregates. These compartments were manually annotated as separate regions of interest on the corresponding whole-slide images.

### Chromogenic immunohistochemistry

Chromogenic immunohistochemistry was performed on formalin-fixed, paraffin-embedded tissue sections using a Dako Autostainer (Agilent Technologies). Slides were deparaffinized and subjected to heat-induced epitope retrieval in high-pH Target Retrieval Solution for 20 minutes. Endogenous peroxidase activity was blocked using an Agilent peroxidase-blocking reagent. Sections were incubated with primary antibodies for 60 minutes at the following dilutions: CD3, 1:400; CD8, 1:600; PD1, 1:600; TIGIT, 1:1,000; TIM3, 1:200; LAG3, 1:500; and GZMB, 1:7,500. A rabbit linker was included in the GZMB detection protocol. Following primary-antibody incubation, staining was detected using Agilent reagents and visualized with 3,3′-diaminobenzidine (DAB) or Magenta chromogen (GV92511-2, Agilent Technologies), as appropriate. Slides were counterstained with Hematoxylin QS (Vector Laboratories; H-3401-500), dehydrated, cleared, and coverslipped.

Primary antibodies included CD3 (Dako/Agilent; A0452), CD8 (Dako/Agilent; M7103), PD1 (Abcam; ab275349), TIGIT (BLR047F, Fortis Life Sciences), TIM3 (Cell Signaling Technology; 45208), GZMB(Novus biological, NBP2-59678-0.1mg) and LAG3 (LifeSpan BioSciences; LS-C344932). TIGIT staining was independently validated using a second anti-TIGIT antibody (Abcam; ab243903).

For dual chromogenic staining, the immune-regulatory or functional marker was visualized with DAB and the corresponding T-cell lineage marker with Magenta. The combinations included PD1/CD3, TIGIT/CD3, TIM3/CD3, LAG3/CD3, and GZMB/CD8. PD1, TIGIT, TIM3, LAG3, and GZMB were developed with DAB, whereas CD3 or CD8 was developed with Magenta. The rabbit or mouse linker used in some staining sequence was selected according to the host species of the corresponding primary antibody. Dual chromogenic immunohistochemistry was performed by sequential staining and detection.

### Multiplex immunofluorescence

Multiplex immunofluorescence was performed on formalin-fixed, paraffin-embedded tissue sections using Opal tyramide signal-amplification reagents (Akoya Biosciences). Sequential cycles of primary-antibody incubation, horseradish peroxidase–mediated detection, fluorophore deposition, and heat-mediated antibody removal were performed.

For the CD8-exhaustion panel, TIM3 (R&D Systems; goat polyclonal; AF2365) was used at 1:750 in Da Vinci Green Diluent. Rabbit anti-goat IgG H&L–HRP (Abcam; ab97100), diluted 1:750 in Da Vinci Green Diluent, was used as the secondary antibody, followed by Opal 620 at 1:150. CD8 (Dako/Agilent; clone 144B; M7103) was used at 1:2,500 and detected with Opal 480 at 1:50. LAG3 (LifeSpan BioSciences; clone 17B4; LS-C341745) was used at 1:3,000 in Da Vinci Green Diluent and detected with Opal 570 at 1:50. PD1 (Cell Signaling Technology; clone EH33; 43248) was used at 1:2,500 and detected with Opal 690 at 1:100. CD163 (Leica/Novocastra; clone 10D6; NCL-L-CD163) was used at 1:200 and detected using TSA-DIG at 1:100, followed by Opal 780 at 1:25. C1Q was detected using a mouse monoclonal antibody against C1QA (Abcam; clone C1QA/2956; ab268120) linked to Opal 520.

For the CD4/Treg/B-cell panel, CD21 (Leica/Novocastra; clone 2G9; NCL-L-CD21-2G9) was detected with Opal 480, CD20 (Dako/Agilent; clone L26; M0755) with Opal 520, CD3 (Dako/Agilent; clone F7.2.38; M7254) with Opal 570, CD4 (Dako/Agilent; clone 4B12; M7310) with Opal 620, PD1 (Cell Signaling Technology; clone EH33; 43248) with Opal 690, and FOXP3 (Cell Signaling Technology; clone D2W8E; 98377) with Opal 780.

The Opal reagents used across the multiplex panels included Opal 480 (FP1500001KT), Opal 520 (FP1487001KT), Opal 570 (FP1488001KT), Opal 620 (FP1495001KT), Opal 690 (FP1497001KT), and Opal 780/TSA-DIG (FP1501001KT). An Opal 3-Plex Detection Kit (Akoya Biosciences; NEL820001KT) was also used. After completion of all staining cycles, slides were counterstained with DAPI and mounted using ProLong Diamond Antifade Mountant (Thermo Fisher Scientific; P36961).

### Slide scanning and digital image analysis

Chromogenic brightfield immunohistochemistry slides were digitized at 20× magnification using a MoticEasyScan One scanner with the Advanced software package (Motic; ES-ONE-20XBUNDLE). Multiplex immunofluorescence slides were digitized using a Leica Biosystems fluorescence whole-slide scanner. Whole-slide images were analyzed using HALO version 4.2 (Indica Labs). Tumor, benign tissue, and lymphoid aggregates were manually annotated as separate regions of interest, and areas containing tissue-processing, scanning, or staining artifacts were excluded from the analysis.

Marker-specific HALO algorithms were used for cell segmentation, phenotyping, and quantification of positive-cell densities and relative immune-cell frequencies For spatial analysis, annotated tissue regions were subdivided into 1-mm^2^ tiles to evaluate the distribution and intratumoral heterogeneity of immune-cell infiltration. Multiplex cellular phenotypes were defined according to the expression and coexpression of the markers included in each panel. Lymphoid aggregates were identified histologically, manually annotated as separate regions of interest, and compared with matched tumor and benign regions. A high density threshold of > 3000 CD3-positive cells/mm^2^ was used to define lymphoid aggregates.

Tumors were classified according to overall CD3-positive T-cell density as Hot (>250 CD3-positive cells/mm^2^) or Cold (≤250 CD3-positive cells/mm^2^). Where direct CD4 staining was unavailable in the chromogenic IHC dataset, CD4-positive T-cell density was estimated from the difference between CD3-positive and CD8-positive cell densities. The GZMB-associated cytotoxic-skewing ratio was calculated as the number of GZMB-positive cells divided by the number of GZMB-negative CD8-positive cells.

### CIBERSORTx immune-cell deconvolution

Immune-cell composition was estimated from bulk RNA-sequencing TPM data using CIBERSORTx in the Impute Cell Fractions module with the LM22 leukocyte signature matrix. The 19 immune-infiltrated prostate tumor punches and 193 TCGA high-Gleason prostate cancers were analyzed together. B-mode batch correction was applied, quantile normalization was disabled because the input consisted of RNA-sequencing data, and significance was assessed using 100 permutations. Both relative immune-cell fractions and absolute immune-abundance scores were generated. Differences between cohorts were evaluated using two-sided Mann–Whitney U tests, with Benjamini–Hochberg correction across the 22 LM22 cell populations.

### RNA isolation, library preparation, and sequencing

Formalin-fixed, paraffin-embedded prostatectomy specimens were histologically reviewed, and immune-infiltrated tumor regions were identified on corresponding tissue sections. A 2-mm core was obtained from each selected region, and RNA was isolated using a Qiagen kit (Qiagen; catalog no. 73604). Libraries were prepared from 100 ng of RNA using the NEBNext Ultra II Directional RNA Library Prep Kit for Illumina (New England Biolabs; E7760), following ribosomal RNA depletion with the NEBNext rRNA Depletion Kit v2 (New England Biolabs; E7400).

Libraries were sequenced on an Illumina NovaSeq platform using an S4 flow cell and a 150-bp paired-end protocol (2 × 150 bp). Sequencing reads were aligned to the human reference genome hg19 and quantified at the gene and transcript levels using an RSEM-based workflow. Twenty immune-infiltrated tumor-punch samples were sequenced. One sample, PCA1, was excluded from downstream analyses because library-quality-control assessment showed an atypical read-distribution profile, including a reduced proportion of reads mapping to coding-sequence exons relative to the remaining samples. The final analysis therefore included 19 tumor punches.

### Comparison with TCGA prostate cancers

RNA-sequencing data from the 19 immune-infiltrated prostate tumor punches were compared with data from 193 primary prostate cancers with Gleason score ≥8 in The Cancer Genome Atlas prostate adenocarcinoma cohort (TCGA-PRAD). The TCGA and tumor-punch datasets were quantified using the same RSEM-based processing workflow to minimize computational differences between cohorts. Gene-level expected counts and transcripts-per-million values were harmonized by gene symbol, and only genes represented in both datasets were retained.

TPM distributions were visualized using violin plots and compared using two-sided Mann–Whitney U tests. For the transcriptome-wide comparisons shown in Figures 4B–F, multiple-testing correction was performed using the Benjamini–Hochberg method across 24,075 analyzed genes, with adjusted P values reported as false-discovery rates. NT5E is included in Figure 4C as an exploratory focused comparison of the CD39–CD73–A2A adenosine-regulatory axis; the displayed two-sided Mann–Whitney U-test P value is nominal, and an FDR was not calculated for this focused comparison. The analyses examined prespecified gene groups related to T-cell lineage and receptor signaling, costimulation, immune checkpoints and adenosine signaling, cytotoxic and natural-killer-cell function, interferon signaling, MHC class I antigen presentation, myeloid and macrophage programs, stromal and vascular components, prostate epithelial differentiation, and proliferation.

### Gene-set enrichment analysis

Gene-set enrichment analysis was used to compare immune-infiltrated tumor punches with high-Gleason TCGA tumors using the matched RSEM-derived expression dataset described above. The analysis evaluated Gene Ontology cellular-component, Reactome, and Hallmark gene sets. Figure S9C shows the proteasome-complex, antigen-processing and cross-presentation, MHC class I antigen-presentation, and interferon-α-response gene sets; false-discovery-rate q values are reported for each enrichment plot.

### Statistical analysis

Relationships between continuous immune measurements were assessed using correlation analysis. Pearson correlation coefficients were used for the reported associations between overall CD3-positive T-cell density and CD8/CD3, TIGIT/CD3, or PD1/CD3 measurements. Matched tissue compartments were compared using two-sided Wilcoxon signed-rank tests. Comparisons between Hot and Cold tumors were performed using two-sided Mann–Whitney U tests. When an overall comparison across three matched compartments was performed, a Friedman test was used, followed by paired Wilcoxon signed-rank tests as appropriate.

All statistical tests were two-sided. Unadjusted p values below 0.05 are described as nominal associations unless the endpoint was prespecified or a stated multiple-testing correction was applied; adjusted significance was assessed using the correction procedure and family defined for each analysis. Cases with missing measurements were excluded only from analyses requiring the unavailable variable.

## Results

### Subset of PCa have high T cell density

We previously reported that a subset of primary PCa had substantial focal CD8 T cell infiltration [6]. To further address whether this reflects a distinct subset of primary PCa that may be eliciting T cells responses, we quantified T cells in a larger series of intermediate to high-risk primary PCa. These included cases that had multiple foci of immune infiltration that were readily apparent on H&E staining to those with sparse immune infiltrates (**Figure S1A**). We used anti-CD3 staining to initially quantify T cell density across entire slides (**Figure S1B**), and arranged the tumors based on overall T cell density (CD3+ cells/mm^2^) (**Figure 1A**).

**Figure 1.**
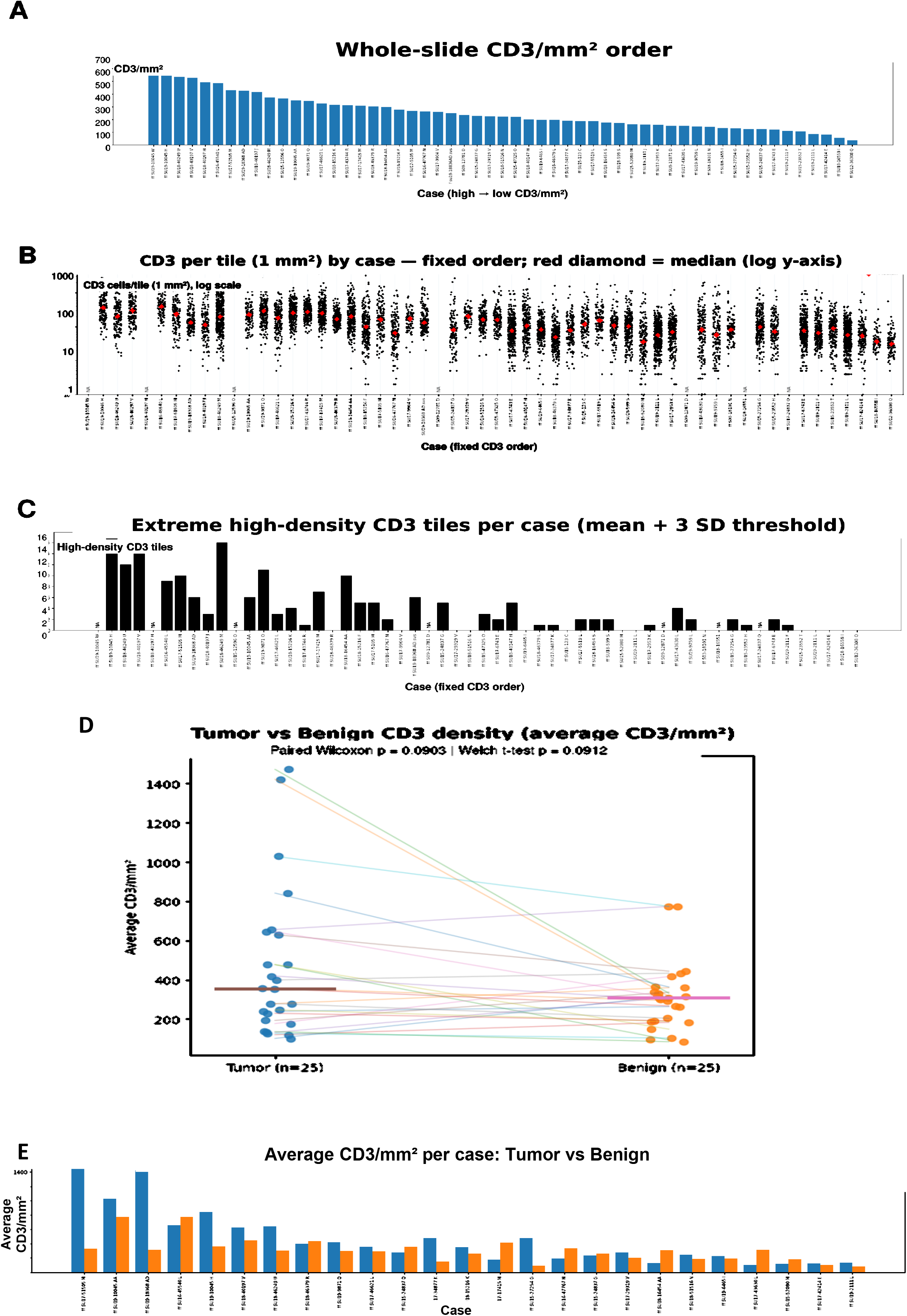
Whole-slide and spatial quantification of CD3⁺ T-cell infiltration in untreated primary prostate cancer. (A) Mean whole-slide CD3⁺ cell density for each case, ordered from highest to lowest. (B) Distribution of CD3⁺ cell density across 1-mm^2^ tiles within each case; each point represents one tile and red diamonds indicate case medians. Cases are ordered as in panel A. (C) Number of extreme high-density CD3⁺ tiles per case, defined as tiles with CD3⁺ density greater than three standard deviations above the overall tile mean. (D) Paired comparison of average CD3⁺ cell density in matched tumor and benign regions. (E) Average CD3⁺ cell density in matched tumor and benign regions, ordered by tumor CD3⁺ density. Cases with unavailable measurements are indicated as not available.

To evaluate the heterogeneity of T cell infiltration at higher spatial resolution, we divided each slide into 1 mm^2^ tiles and quantified the number of CD3⁺ cells in each 1 mm^2^ tile (**Figure 1B, Figure S2**). Cases were ordered as in figure 1A according to their overall T cell density. Across all cases, the number of T cells per tile ranged from 0 to 5,852, with a median of 122 T cells /mm^2^ and a mean of 213 T cells/mm^2^. We also quantified the number of 1 mm^2^ tiles with T cell density greater than 3 SD above the mean (**Figure 1C**). This analysis indicates that increased focal T cell accumulation is correlated with increased overall T cell density, although smaller numbers of focal T cell hot spots could be found in tumors with lower overall T cell density.

We next compared T cell density between matched tumor and benign compartments. Tumor areas showed increased T cell density and greater heterogeneity in T cell infiltration than benign regions (**Figure 1D**). Notably, cases with higher T cell density in their tumor compartments had higher overall T cell density, and in most cases had lower density in their corresponding benign areas, with the density in the benign areas being only modestly greater than in the tumors with lower overall T cell density (**Figure 1E**). Together, these results show that increased T cell density is driven primarily by T cell accumulation in tumor foci.

### Increased T cell density is associated with decreased CD8/CD4 T cell ratio

We next used dual CD8/CD3 IHC to similarly characterize CD8 T cells overall density and spatial distribution (**Figure S3**). Cases were again ordered based on the above overall T cell density, which showed that higher CD8 T cell density was generally associated with higher overall T cell density (**Figure 2A**). Analysis using 1 mm^2^ grid segmentation then revealed substantial variation in CD8⁺ T cell density across and within cases (**Figure 2B and Figure S4A**).

**Figure 2.**
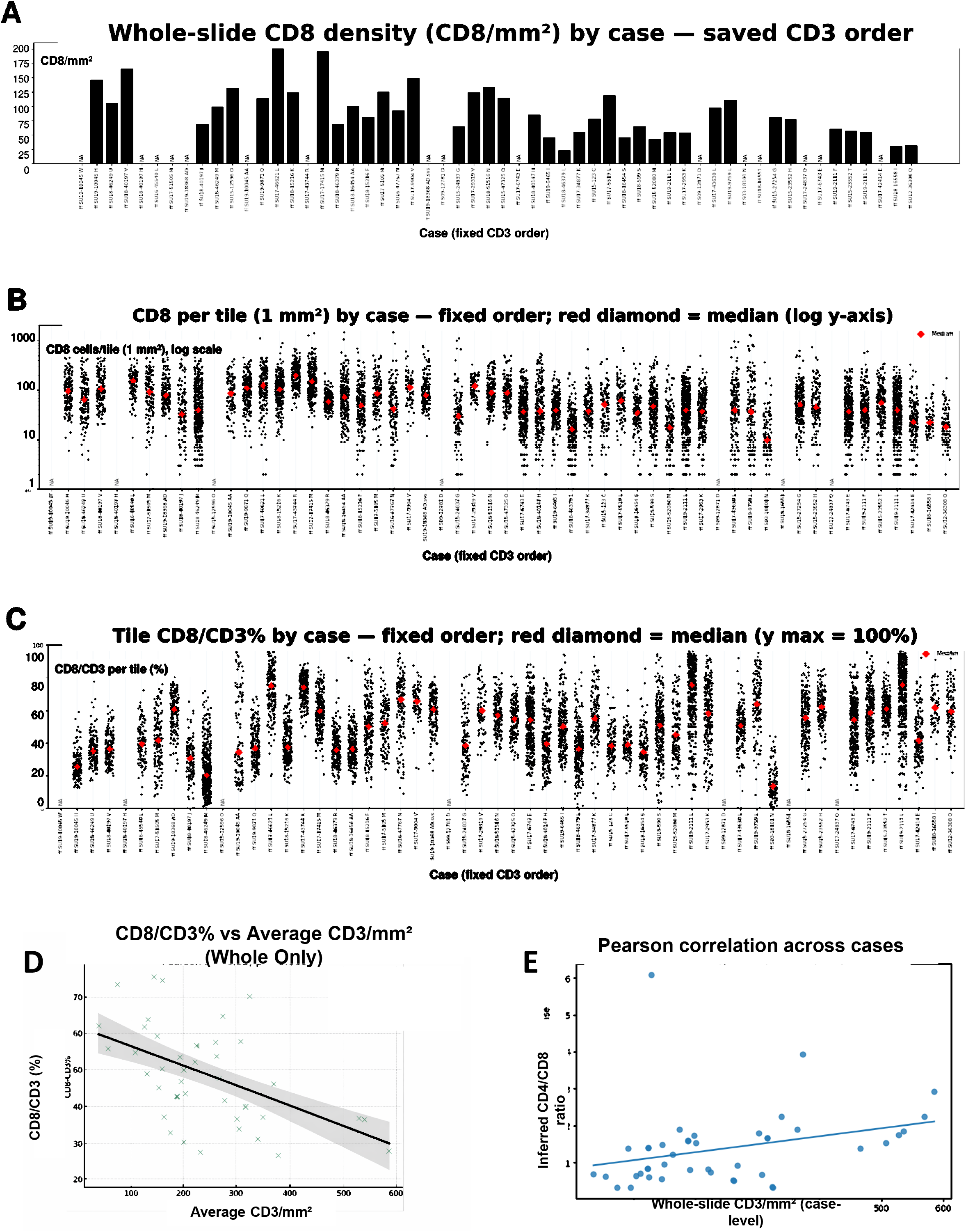
Increasing T-cell infiltration is associated with reduced proportional CD8⁺ T-cell representation. (A) Mean whole-slide CD8⁺ cell density by case, with cases ordered by whole-slide CD3⁺ density. (B) Distribution of CD8⁺ cell density across 1-mm^2^ tiles; each point represents one tile and red diamonds indicate case medians. (C) Tile-level CD8/CD3 ratio across cases; each point represents one 1-mm^2^ tile and red diamonds indicate case medians. (D) Inverse relationship between the case-level CD8/CD3 ratio and average whole-slide CD3⁺ cell density (Pearson r = −0.53, p < 0.001); the line shows the linear fit and shading shows the 95% confidence interval. (E) Relationship between the inferred CD4/CD8 ratio and whole-slide CD3⁺ cell density (Pearson r = 0.304, p = 0.0472; n = 43). CD4⁺ cell abundance was inferred as CD3⁺ minus CD8⁺ cells.

We next examined the CD8/CD3 ratio at the tile level across all cases. For each 1 mm^2^ tile, the CD8/CD3 ratio was calculated and plotted as a single point, while the median CD8/CD3 ratio per case was overlaid as a red diamond, with cases again ordered by descending overall T cell density (**Figure 2C**). Across all 8,127 tiles, CD8/CD3 values spanned the full range from 0–100% (mean ≈ 44.68 %, median ≈ 42.16 %), indicating that on average roughly half of T cells were CD8⁺, but with substantial variability both within and between cases. Notably, examining all the cases it appeared that the CD8/CD3 ratio was generally lower in tumors that had greater T cell infiltration (based on median T cell density). To assess this relationship between overall T cell density and CD8 T cell composition, we correlated overall T cell density (CD3/mm^2^) with mean CD8/CD3 ratio for each case. This revealed a significant inverse correlation between the two metrics (Pearson r = –0.53, p < 0.001) (**Figure 2D**). This finding indicates that although tumors with higher overall T-cell infiltration have increased CD8 T cells, non-CD8 T cells show greater proportional representation.

Consistent with the above result, analysis of inferred CD4^+^ T cells (derived by subtracting CD8⁺ cells from total CD3⁺ T cells) shows a strong correlation with overall CD3⁺ T-cell density (**Figure S4B**). Assessment of inferred CD4^+^ T cells per 1 mm^2^ tile showed substantial intra-tumoral heterogeneity, with densities ranging from 13 to 260 cells per tile, and an overall mean of ∼140 CD4⁺ T cells per 1 mm^2^ tile across all tiles (**Figure S4C, D**). As expected based on the above CD8^+^ T cell result, there was a significant positive correlation between CD4/CD3 ratio and T cell density (**Figure S4E**). Finally, plotting CD4/CD8 T cell ratio versus CD3 density further shows that increased T cell infiltration is associated with a relative increase in the CD4/CD8 T cell ratio (**Figure 2E**).

In a subset of cases we also examined PD1 expression (**Figure S5A**). The fraction of T cells expressing PD1 was not clearly correlated with T cell density (**Figure S5B**), and graphical analysis confirmed that there was no overall association with CD3 density (**Figure S5C**).

### CD8 and CD4 T cell phenotypes in immune infiltrated foci

We next stratified cases by overall T cell density, defining Hot tumors as >250 CD3-positive cells/mm^2^ and Cold tumors as ≤250 CD3-positive cells/mm^2^ (**Figure S6A**). We then used two multiplex IF panels to assess whether this CD3 density-based classification captured differences in T cell, B cell, or macrophage phenotypes. The first panel targeted CD8, PD1, LAG3, TIM3, CD163, and C1Q **(Figure S6B, C).** Consistent with the above IHC analysis, the fraction of CD8 T cells that were PD1⁺ was similar between Hot and Cold tumors (**Figure 3A**). We next focused on the PD1⁺CD8⁺ population and quantified expression of TIM3 and LAG3. Hot tumors had higher fractions of TIM3⁺ and LAG3⁺ subsets within PD1⁺CD8⁺ cells (**Figure S6D-F**), and conversely had a lower fraction that were negative for both TIM3 and LAG3 (**Figure S6G**), although there was a broad range in the Hot tumors and the differences did not reach statistical significance. Based on these data we also carried out IHC studies to assess TIM3 and LAG3 in additional cases **(Figure S7A, B).** Consistent with above multiplex IF, there was a trend toward higher TIM3 and LAG3 expression in T cells in Hot versus Cold tumors (**Figure S7C, D**). Quantitatively, median TIM3/CD3 was 3.65% (IQR, 2.01–4.82; n = 18) in Hot tumors versus 2.97% (IQR, 2.47–3.67; n = 10) in Cold tumors (two-sided Mann–Whitney U P = 0.8667). Median LAG3/CD3 was 7.16% (IQR, 4.76–10.73; n = 18) in Hot tumors versus 4.45% (IQR, 2.82–6.07; n = 11) in Cold tumors (two-sided Mann–Whitney U P = 0.3119).

**Figure 3.**
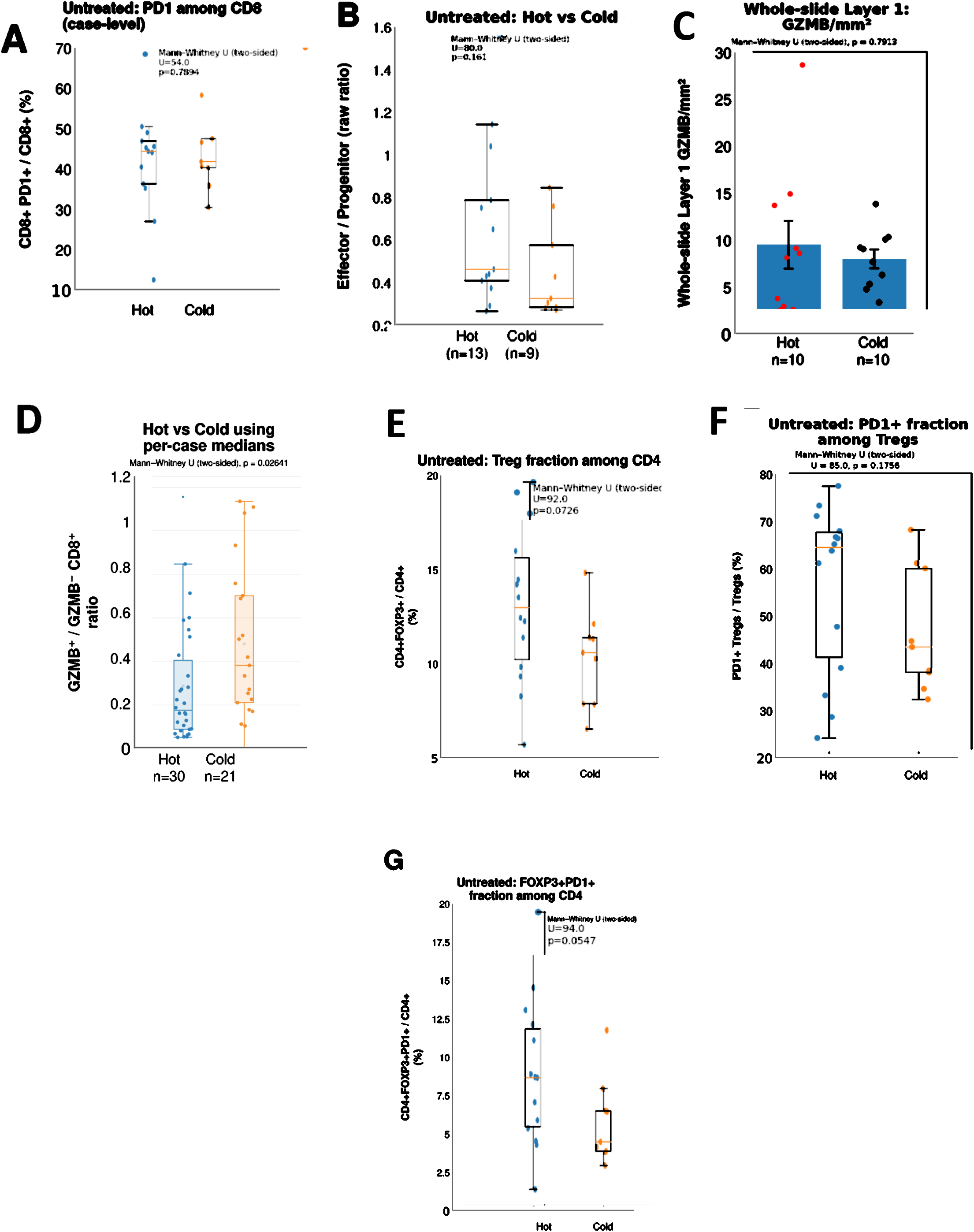
T-cell phenotype, GZMB-associated cytotoxic skewing, and regulatory T-cell features in Hot and Cold tumors. Tumors were classified as Hot or Cold using a whole-slide CD3⁺ density threshold of 250 cells/mm^2^. (A) Percentage of CD8⁺ T cells expressing PD1. (B) Ratio of effector/exhausted to progenitor-like cells within the PD1⁺CD8⁺ population; effector/exhausted cells were defined as TIM3⁺ and/or LAG3⁺, and progenitor-like cells as TIM3⁻LAG3⁻. (C) Whole-slide GZMB⁺ cell density. (D) GZMB abundance relative to GZMB-negative CD8⁺ cells; the ratio was higher in Cold tumors (two-sided Mann–Whitney U test, p = 0.0264). (E) FOXP3⁺ regulatory T cells as a percentage of CD4⁺ T cells. (F) PD1⁺ cells as a percentage of FOXP3⁺CD4⁺ regulatory T cells. (G) FOXP3⁺PD1⁺ cells as a percentage of CD4⁺ T cells. Points represent individual tumors; box plots show the median and interquartile range, and bar plots show the mean with error bars as displayed. Hot-versus-Cold comparisons used two-sided Mann–Whitney U tests.

Notably, we had reported previously in an examination of focally immune infiltrated tumors that many of the CD8^+^ PD1^+^ TIM3⁻ LAG3⁻ cells were expressing TCF1, consistent with stem-like/progenitor T cells [6]. Therefore, we computed a per-case exhausted versus progenitor cell ratio within the PD1⁺CD8⁺ T cell subset, defining exhausted cells as TIM3⁺ and/or LAG3⁺ and progenitor cells as negative for TIM3 and LAG3. This ratio trended higher in the Hot tumors, but did not reach statistical significance **(Figure 3B).** These results indicate a directional shift from progenitor-like toward more recently antigen-experienced phenotypes in Hot tumors, although the difference did not reach statistical significance.

As GZMB was not in the multiplex IF panel, we next asked whether the CD8⁺ T cells present in the Hot versus Cold tumors showed evidence of enhanced cytotoxic effector differentiation. To address this, we used dual chromogenic IHC to co-quantify GZMB-expressing cells and CD8⁺ T cells (**Figure S8**). Overall the density of GZMB+ cells was comparable in the Hot versus Cold tumors (**Figure 3C**). However, the ratio of GZMB positive cells versus GZMB negative CD8 cells was significantly higher in the Cold tumors (**Figure 3D**). Notably, as we could not assess CD8 staining in cells staining positive for GZMB, it is possible that some GZMB positive cells were NK cells. Nonetheless, together these data indicate that while an increased fraction CD8^+^ stem-like/progenitor T cells may become activated in the Hot tumors, their cytotoxic effector function is reduced in the tumor microenvironment.

The second multiplex IF panel targeted CD3, CD4, FOXP3, PD1, CD20, and CD21. Although not statistically significant, the Hot tumors had a higher fraction of CD4 T cells expressing FOXP3 (**Figure 3E**) and a higher fraction of these CD4^+^FOXP3^+^ cells had PD1 expression (**Figure 3F**). Consistent with these findings, the fraction of CD4 T cells expressing both FOXP3 and PD1 was greater in the Hot tumors, and this difference approached statistical significance (**Figure 3G**). In conjunction with the shift towards higher CD4/CD8 T cell ratios in the Hot tumors, these patterns are consistent with more T cell infiltrated tumors engaging compensatory immunosuppressive programs. Finally, the proportion of CD163⁺ macrophages that co-expressed C1Q was comparable between Hot and Cold tumors (**Figure S7E).**

### RNA-seq analysis of Hot tumors

We took 2 mm punches from immune infiltrated areas in Hot tumor cases and carried out RNA-seq. We first applied CIBERSORTx with the LM22 reference signature to 19 evaluable cases and compared the inferred cell fractions with those from 193 TCGA prostate cancers with Gleason score ≥8. Eleven of the 22 inferred immune populations differed significantly after multiple-testing correction (**Figure 4A, Figure S9A, B**). The Hot punches had significantly higher median fractions of resting memory CD4⁺ T cells (30.28% versus 19.38%; FDR = 7.7 × 10⁻⁶), resting NK cells (3.02% versus 0%; FDR = 2.5 × 10⁻⁶), and monocytes (4.26% versus 2.50%; FDR = 0.0065) than TCGA tumors. Conversely, the relative fractions of M0, M1, and M2 macrophages, follicular helper T cells, and regulatory T cells were lower in the Hot punches (all FDR < 0.05). Notably, the inferred CD8⁺ T-cell fraction was also lower (9.09% versus 12.20%), although this difference narrowly missed significance after FDR correction (P = 0.0287; FDR = 0.0527) (**Figure S9A, B**). This result is consistent with the IHC analysis in showing a relative enrichment of CD4^+^ versus CD8^+^ T cells in the Hot tumors.

**Figure 4.**
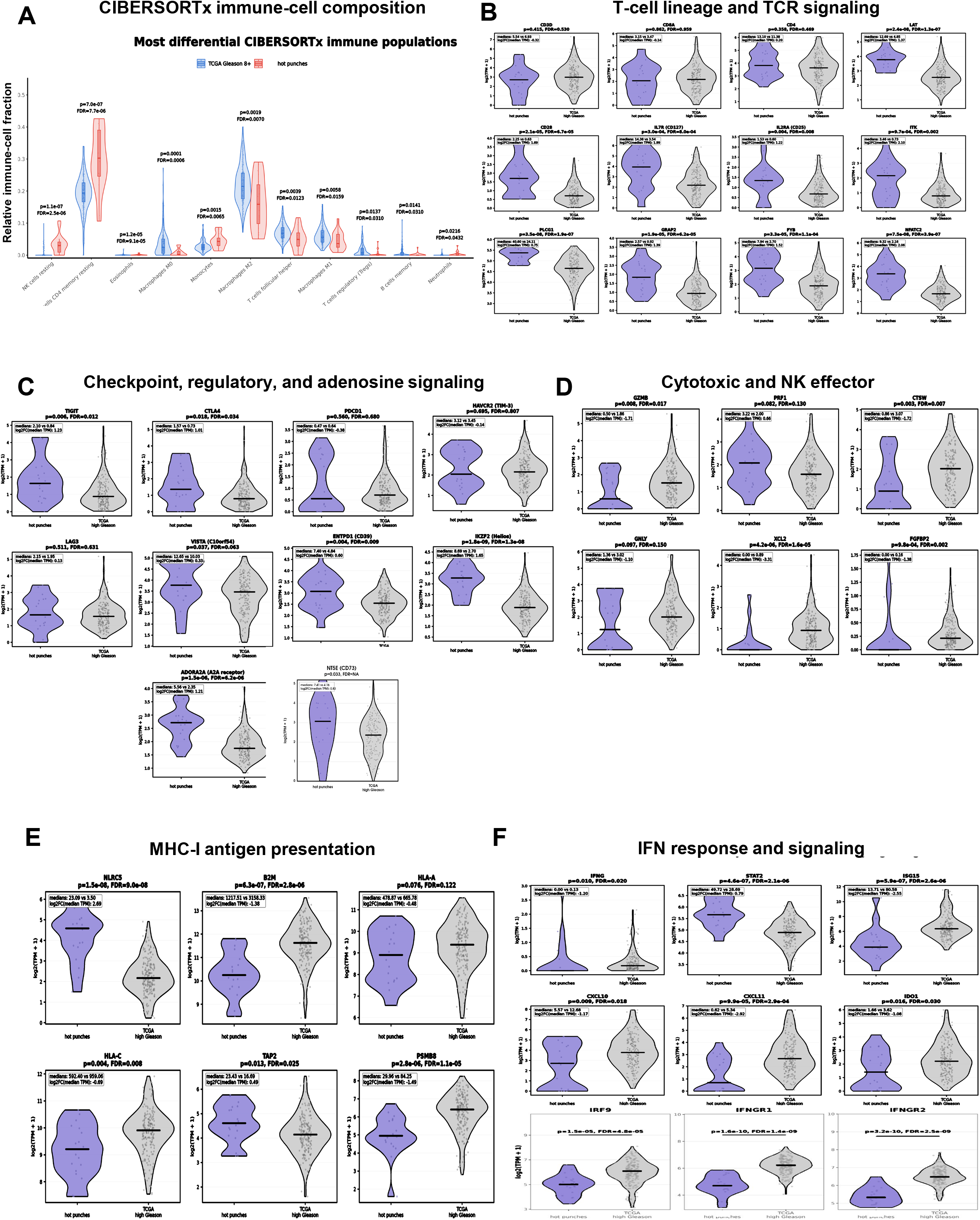
Immune-cell composition and immune-regulatory transcriptional programs in Hot punches. Nineteen evaluable Hot punches were compared with 193 TCGA primary prostate cancers with Gleason score ≥8. (A) CIBERSORTx LM22 relative fractions for the most differentially represented immune-cell populations. Transcript-per-million comparisons are organized by biological group: (B) T-cell lineage and T-cell receptor signaling; (C) immune-checkpoint, immune-regulatory, and adenosine-signaling genes; (D) cytotoxic T-cell and natural-killer-cell effectors; (E) MHC class I antigen-presentation genes; and (F) interferon-response and inflammatory-signaling genes together with IFNGR1, IFNGR2, and IRF9. Benjamini–Hochberg-adjusted FDR values are shown for the genome-wide comparisons. NT5E is included as an exploratory focused comparison with a nominal two-sided Mann–Whitney U P value and no calculated FDR. Points represent individual samples and horizontal lines indicate the displayed distribution summaries.

We next evaluated overall transcript abundance across predefined T-cell-lineage, T-cell-signaling, immune-regulatory, cytotoxic, antigen-presentation, interferon-response, myeloid, stromal, epithelial, and proliferation-related gene groups. Despite their selection from regions with dense immune infiltration, the Hot punches did not show a generalized increase in T-cell-lineage transcripts (**Figure 4B**). Expression of CD3D and CD8A was comparable between the cohorts, and CD4 was also not significantly increased in the Hot punches (median, 13.10 versus 11.38 TPM; FDR = 0.469). The basis for this is not clear, but could possibly reflect relative downregulation of mRNA levels for these antigen recognition proteins in the Hot tumor foci.

In contrast, the Hot punches showed enrichment of selected T-cell receptor-signaling and activation-associated genes (**Figure 4B**). These included LAT, a scaffold that organizes signaling immediately downstream of the T-cell receptor; ITK, a T-cell kinase that propagates receptor signals; PLCG1, which initiates calcium- and diacylglycerol-dependent signaling; and the adaptor proteins GRAP2 and FYB, which couple T-cell receptor signaling to adhesion and cytoskeletal responses. The downstream transcription factor NFATC2 was also increased. Together with increased CD28, IL7R/CD127, and IL2RA/CD25, these findings suggest enhanced signaling or activation within the infiltrating T-cell population.

The immunoregulatory profile was selective rather than reflecting uniform upregulation of all inhibitory receptors (**Figure 4C**). TIGIT expression was significantly higher in the Hot punches than in TCGA tumors (median, 2.10 versus 0.84 TPM; FDR = 0.012). There was also increased ENTPD1/CD39 (median, 7.40 versus 4.84 TPM; FDR = 0.009), an ectonucleotidase that converts extracellular ATP and ADP to AMP, which can subsequently be converted to adenosine by NT5E/CD73. ADORA2A, which encodes the A2A receptor, was also increased (median, 5.56 versus 2.35 TPM; transcriptome-wide FDR = 6.2 × 10⁻⁶). A focused visualization of the CD39–CD73–A2A adenosine-regulatory axis showed higher expression of all three components in the Hot punches (**Figure 4C**). Adenosine is the ligand for the A2A receptor, and engagement of this receptor suppresses T-cell activation and effector function. Together, these findings support a transcriptionally enriched candidate CD39–CD73–adenosine–A2A immunoregulatory axis.

CTLA4 was increased (median, 1.57 versus 0.73 TPM; FDR = 0.034), as was IKZF2/Helios (median, 8.69 versus 2.70 TPM; FDR = 1.3 × 10⁻⁸), a transcription factor associated with regulatory and chronically stimulated T-cell states (**Figure 4C**). VISTA/C10orf54, an inhibitory checkpoint expressed prominently by myeloid cells and subsets of T cells, showed a nominal increase (median, 12.65 versus 10.03 TPM; P = 0.037) that did not remain significant after transcriptome-wide correction (FDR = 0.063). In contrast, PDCD1/PD-1, HAVCR2/TIM-3, and LAG3 were unchanged. Thus, the regulatory state was not characterized by coordinated elevation of every exhaustion-associated receptor, but particularly by TIGIT-, CTLA4-, CD39–A2A-adenosine-, and Helios-associated programs, with VISTA showing a nonsignificant upward trend (**Figure 4C**).

This regulatory profile was not accompanied by a broadly enhanced cytotoxic-effector program. Consistent with the IHC data, GZMB expression was significantly lower in the Hot punches than in TCGA tumors (median, 0.50 versus 1.86 TPM; FDR = 0.017) (**Figure 4D**). CTSW, a protease enriched in cytotoxic lymphocytes; XCL2, a chemokine produced by activated cytotoxic T and NK cells; and FGFBP2, a marker of highly differentiated cytotoxic T and NK populations, were also reduced. GNLY, which encodes the NK- and cytotoxic T-cell effector protein granulysin, was numerically lower but was not significant after multiple-comparison correction. PRF1, encoding perforin, was numerically higher but also did not remain significant. Thus, the immune-infiltrated foci lacked a coordinated granzyme- and NK-associated effector program despite retaining selected components of the cytolytic machinery.

Antigen-presentation genes also showed a discordant pattern (**Figure 4E**). NLRC5, the principal transcriptional activator of MHC class I genes, was markedly increased in the Hot punches (median, 23.09 versus 3.50 TPM; FDR = 9.0 × 10⁻⁸). TAP2, which transports proteasome-derived peptides into the endoplasmic reticulum for loading onto MHC class I molecules, was also increased (median, 23.43 versus 16.69 TPM; FDR = 0.025). In contrast, B2M, HLA-C, and PSMB8 were significantly reduced. PSMB8 encodes an inducible immunoproteasome subunit that contributes to the production of peptides for MHC class I presentation. HLA-A was numerically lower but was not significant after multiple-comparison correction.

Interferon-associated genes showed a mixed pattern (**Figure 4F**). STAT2, a mediator of type I interferon signaling, was increased, whereas IFNG, ISG15, CXCL10, CXCL11, IDO1, IRF9, IFNGR1, and IFNGR2 were reduced. IRF9, IFNGR1, and IFNGR2 remained significantly lower after multiple-testing correction (FDR = 4.8 × 10⁻⁵, 1.4 × 10⁻⁹, and 2.5 × 10⁻⁹, respectively) (**Figure 4F**). CXCL10 and CXCL11 encode interferon-inducible chemokines that recruit activated CXCR3⁺ lymphocytes, whereas IDO1 encodes an interferon-inducible enzyme that suppresses T-cell responses through tryptophan metabolism. Consistent with this discordant gene-level pattern, gene-set enrichment analysis showed reduced enrichment of proteasome-complex, antigen-processing and cross-presentation, MHC class I antigen-presentation, and interferon-α-response pathways in Hot punches relative to high-Gleason TCGA tumors (**Figure S9C**). Together, these findings indicate partial or uncoupled engagement of interferon signaling and antigen-presentation pathways rather than a uniformly activated interferon–MHC class I program.

The Hot punches also exhibited a compositionally complex myeloid microenvironment (**Figure S10A**). CD163, ITGAM/CD11b, MRC1/CD206, and CSF1R were increased, supporting enrichment of selected monocyte- and macrophage-associated populations. CD163 and MRC1 are frequently associated with scavenging and tissue-remodeling macrophage states, whereas CSF1R is a major regulator of monocyte and macrophage survival. However, FCER1G and AIF1 were reduced, indicating that the myeloid program was not uniformly increased. Notably, SPP1 was markedly lower in the Hot punches than in TCGA tumors (median, 2.90 versus 19.61 TPM; transcriptome-wide FDR = 2.5 × 10⁻⁸). SPP1 encodes osteopontin and is commonly used to identify an SPP1-positive tumor-associated macrophage population associated with extracellular-matrix remodeling, angiogenesis, and tumor progression. Therefore, the increased expression of CD163, MRC1, ITGAM, and CSF1R did not reflect generalized enrichment of all macrophage states and specifically was not accompanied by an SPP1-positive macrophage program.

Stromal genes, including COL1A1, COL1A2, COL3A1, and PDGFRB, were increased, indicating extracellular-matrix and perivascular stromal enrichment (**Figure S10B**). In contrast, proliferation-associated genes, including PCNA, TOP2A, CDK1, CCNB1, CCNB2, and UBE2C, were consistently reduced, suggesting lower proliferative tumor-cell content or activity within the selected immune-infiltrated regions (**Figure S10C**). Finally, although AR mRNA was increased in the Hot tumors, other luminal epithelial genes (EPCAM, KRT8, KLK3, FOXA1, and HOXB13) were decreased (**Figure S10D**).

### TIGIT expression is increased in T cell infiltrated tumors

Based on the above RNA-seq data, we evaluated TIGIT by IHC (**Figure S11A**). The density of TIGIT-expressing cells was highly variable, but was generally increased in tumors with higher overall T cell density, consistent with the RNA-seq data (**Figure 5A**). The TIGIT/CD3 ratio appeared to be less associated with T cell density (**Figure 5B**). However, plotting TIGIT/CD3 ratio versus overall CD3 density showed a nominally significant positive association, indicating that an increased fraction of T cells in immune infiltrated tumors are expressing TIGIT (**Figure 5C**).

**Figure 5.**
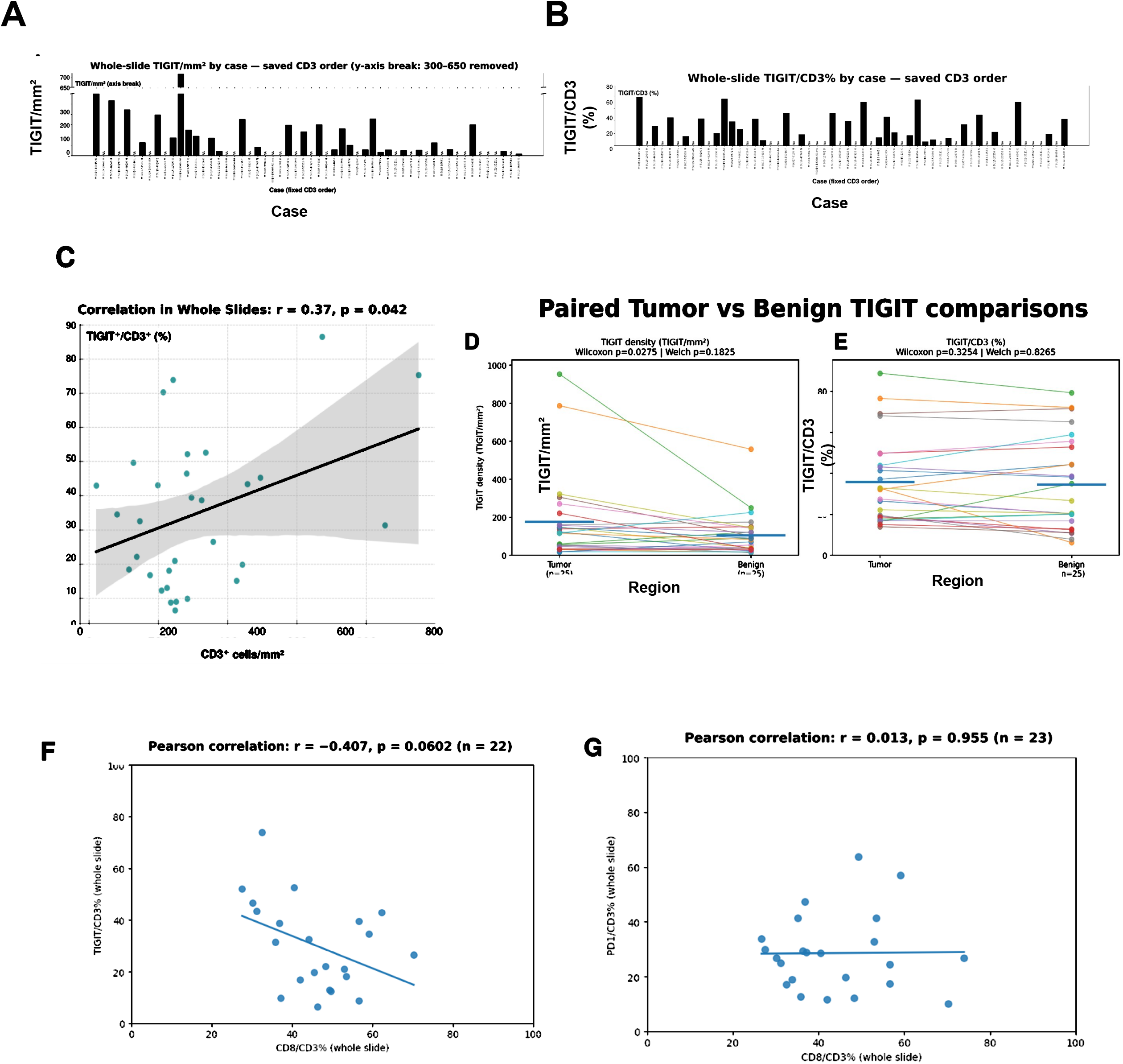
TIGIT expression increases with T-cell infiltration and is inversely associated with CD8 proportional representation. (A) Whole-slide TIGIT⁺ cell density by case, ordered by whole-slide CD3⁺ density. (B) Whole-slide TIGIT/CD3 ratio by case in the same order. (C) Association between TIGIT/CD3 and whole-slide CD3⁺ density (Pearson r = 0.37, p = 0.042); the line shows the linear fit and shading shows the 95% confidence interval. (D) Paired comparison of TIGIT⁺ cell density in matched tumor and benign regions. (E) Paired comparison of TIGIT/CD3 in matched tumor and benign regions. Matched comparisons used two-sided Wilcoxon signed-rank tests. (F) Inverse relationship between TIGIT/CD3 and CD8/CD3 (Pearson r = −0.407, p = 0.0602; n = 22). (G) Relationship between PD1/CD3 and CD8/CD3 (Pearson r = 0.013, p = 0.955; n = 23).

We next compared TIGIT expression between paired tumor and benign regions. TIGIT^+^ cell density was higher in tumor than benign regions (**Figure 5D**). However, the proportion of T cells that are TIGIT positive was similar between tumor and benign regions (**Figure 5E**). Together these findings indicate that a higher fraction of T cells express TIGIT in the immune infiltrated tumors, and that this increase is in the tumor and benign areas.

We also examined the correlation between the TIGIT/CD3 and the CD8/CD3 ratio, which showed they were negatively correlated (**Figure 5F**). Notably, tumors with lower CD8/CD3 ratios (∼30%) had TIGIT/CD3 ratios over 40%, suggesting TIGIT expression among non-CD8 T cells in these cases. In contrast, there was no correlation between the fraction of T cells expressing PD1 and the CD8/CD3 ratio (**Figure 5G**).

To confirm the robustness of TIGIT immunohistochemical quantification, we validated our findings using a second anti-TIGIT antibody (ab243903, Abcam). Quantitative image analysis revealed a strong concordance between the original antibody and the validation clone (**Figure S11B**).

### Lymphoid aggregates are enriched for non-CD8 T cells and TIGIT

For this analysis, we also annotated lymphoid aggregates based on histological assessment (**Figure S12A**). Most contained predominantly CD3⁺ cells, indicating that they were not established mature tertiary lymphoid structures (TLSs), although some could represent early TLSs (**Figure S12B**). The CD8/CD3 ratio was comparable in tumor and benign regions, but lower in lymphoid aggregates, indicating enrichment for non-CD8 T cells (**Figure 6A**). PD1/CD3 was also similar between tumor and benign regions and was higher in lymphoid aggregates (**Figure 6B, C**). Finally, TIGIT⁺ cells were quantified as a percentage of total CD3⁺ T cells across matched tumor, benign, and lymphoid-aggregate regions. Pairwise Wilcoxon signed-rank tests showed that TIGIT/CD3 was higher in lymphoid aggregates than in benign tissue (p = 6.68 × 10⁻⁶) or tumor (p = 0.0093), whereas benign and tumor regions did not differ significantly (p = 0.235) (**Figure 6D**).

**Figure 6.**
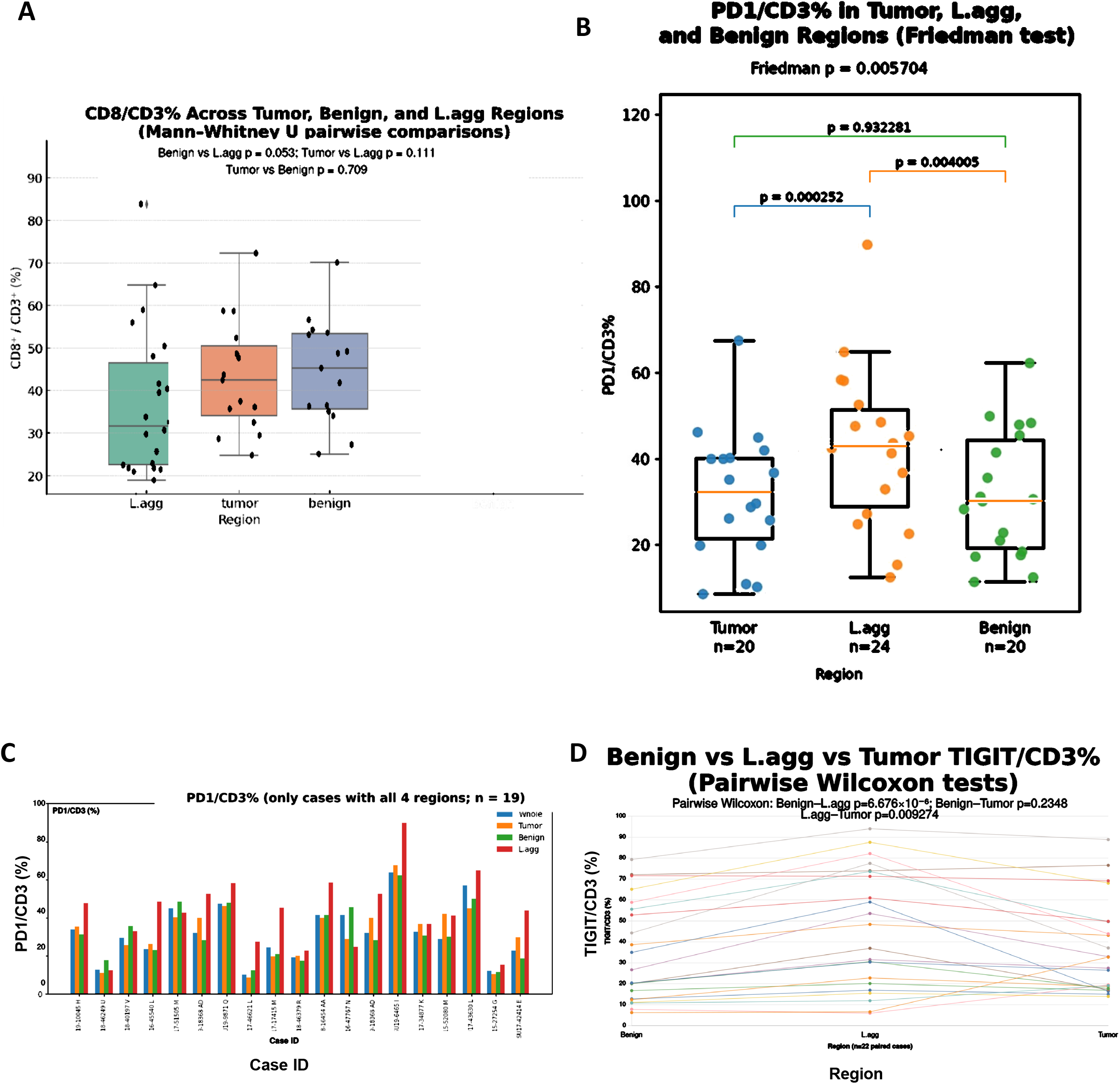
Lymphoid aggregates are enriched for non-CD8 T cells and checkpoint-expressing T cells. Lymphoid aggregates were defined as regions containing >3000 CD3⁺ cells/mm^2^. (A) CD8/CD3 across tumor, benign, and lymphoid-aggregate regions; exploratory pairwise two-sided Mann–Whitney U tests are shown. (B) PD1/CD3 across tumor, lymphoid-aggregate, and benign regions; the overall Friedman test and pairwise comparisons are shown. (C) Case-level PD1/CD3 values across whole-slide, tumor, benign, and lymphoid-aggregate compartments for cases with all four measurements (n = 19). (D) TIGIT/CD3 in matched benign, lymphoid-aggregate, and tumor regions (n = 22); lines connect regions from the same case, and pairwise Wilcoxon signed-rank comparisons are shown. Points or bars with the same color identify measurements from the same case where applicable.

## Discussion

While PCa does not generally appear to elicit T-cell responses, focal immune infiltrates are found in a subset of cases. We selected here a series of untreated primary PCa that had varying degrees of immune infiltration to address whether there may be qualitative differences in tumors that are more heavily versus more sparsely infiltrated. We determined by IHC overall T cell density in these tumors and found that higher T cell density was driven primarily by increased focal tumor infiltration, although cases with higher T cell density in tumor areas also had increased density in benign areas. Notably, while tumors with higher T cell density also had increased CD8 T cell density, we found that higher T cell density was associated with a decrease in the CD8/CD4 T cell ratio. Moreover, higher T cell density was also associated with a decrease in the ratio of GZMB-positive cells to GZMB-negative CD8⁺ T cells. Together these findings suggest that tumors that are stimulating greater immune responses are engaging immunoregulatory mechanisms to suppress CD8 T effector functions.

Further multiplex IF analyses of CD8 T cells in immune infiltrates showed a greater fraction of the PD1 positive cells in the Hot tumors expressed TIM3 and/or LAG3, suggesting they had been recently activated in the tumor microenvironment (TME). In conjunction with the lower GZMB in these Hot cases, this suggests that factors in the TME of the Hot tumors may be impairing the development of effector functions in tumor-reactive CD8 T cells and/or rapidly driving them to exhaustion. One such factor may be regulatory T cells, as the multiplex IF analysis indicated FOXP3 positive CD4 T cells were increased in the Hot tumors and a greater fraction of these were PD1 positive.

Whole transcriptome analysis of immune infiltrated tumor foci suggested further factors in the TME that may be suppressing T cell function. Analysis by CIBERSORTx in comparison with high-Gleason TCGA tumors showed significant enrichment for CD4 resting memory T cells, but not for CD8 T cells, consistent with our IHC data. Resting NK-cell and monocyte fractions were also higher, whereas the relative fractions of follicular-helper T cells, Tregs, and M0, M1, and M2 macrophages were lower, although these differences were not highly significant. Also consistent with the IHC results, expression of GZMB was lower in the infiltrated tumor foci.

The transcriptome analysis also showed selective enrichment of immune checkpoint related genes including TIGIT, CTLA4, ENTPD1/CD39, ADORA2A/A2AR, NT5E/CD73, and IKZF2/Helios. These selective changes support checkpoint adaptation in a chronically stimulated immune environment rather than uniform loss of CD8 T cell effector identity or definitive terminal exhaustion [16–18]. The increased expression of CD39, CD73, and A2AR supports a role for extracellular adenosine in suppressing immune responses [19]. IKZF2 has been implicated in supporting the function of immunosuppressive Tregs [20], and may also be induced in tumor-infiltrating exhausted CD8 T cells [21]. In PCa, a recent study showed IKZF2 expression in Tregs that were increased in prostate after neoadjuvant androgen deprivation therapy [22].

Based on the RNA-seq data we carried out further IHC studies for TIGIT, which confirmed that its increased expression was associated with higher T-cell infiltration. Specifically, both TIGIT-positive cell density and the TIGIT/CD3 ratio increased with T-cell density. This pattern contrasted with the PD1/CD3 ratio, which was not significantly associated with T-cell density. This distinction may be therapeutically relevant because the TIGIT–CD226–PVR axis regulates both effector and regulatory T-cell responses and may remain active in settings in which PD1-directed therapy alone is insufficient [23,24]. Single-cell analysis of localized PCa has likewise identified TIGIT among the inhibitory receptors expressed by CD8⁺ T cells in high-grade tumors [25]. Functionally, TIGIT blockade enhanced the activity of human peripheral-blood NK cells against castration-resistant prostate-cancer cells in preclinical models, supporting TIGIT as a biologically relevant therapeutic target in this disease [26]. Anti-TIGIT antibodies have entered early-phase clinical testing in advanced solid tumors, including tiragolumab alone or with atezolizumab, but prostate-cancer–specific clinical efficacy has not yet been established [27].

Organized lymphoid aggregates, including tertiary lymphoid structures, can facilitate interactions among T cells, B cells, and antigen-presenting cells and thereby support local antigen presentation and lymphocyte activation [28]. In the matched regional analysis, TIGIT/CD3 was significantly higher in lymphoid aggregates than in either matched benign or tumor regions, whereas benign and tumor regions did not differ significantly. PD1/CD3 was likewise enriched within lymphoid aggregates. Persistent antigen exposure within these structures may simultaneously promote checkpoint induction and accumulation of regulatory programs. Their enrichment for both TIGIT and PD1 is therefore compatible with the coexistence of immune activation and compensatory immune restraint. However, most aggregates in this study were predominantly T-cell rich and were not demonstrated to be mature tertiary lymphoid structures. Additional spatial phenotyping will be required to define their organization and determine whether TIGIT and PD1 are coexpressed by the same cells or identify different effector, progenitor-like, follicular-helper, or regulatory T-cell populations.

In conclusion, increasing T-cell infiltration in untreated primary PCa was associated with reduced proportional CD8 representation and reduced GZMB-associated cytotoxic skewing rather than a proportionately stronger effector response. Tissue and bulk analyses demonstrated selected T-cell activation and checkpoint/chronic-stimulation programs, with TIGIT showing a consistent association with immune-rich disease and both TIGIT and PD1 enriched within lymphoid aggregates. The accompanying CD39–CD73–adenosine–A2A, interferon, antigen-presentation, myeloid, and stromal findings further demonstrate that the immune-rich prostate tumor microenvironment is active but compositionally and functionally counter-regulated. T-cell density alone is therefore insufficient to define an effective antitumor immune state. Integrating immune composition, cytotoxic differentiation, checkpoint programs, antigen-presentation capacity, and spatial organization may provide a more informative framework for identifying primary prostate cancers most likely to benefit from immune-targeted strategies.

## Acknowledgments

This work was supported by NIH grants P01 CA163227, R01 CA272934, and P50 CA272390 to SPB. Further support was provided by a Prostate Cancer Foundation Challenge grant to SPB.

## Notes

### Competing Interest Statement

The authors have declared no competing interest.

