## Supplementary Figures for "Higher T-cell density in primary prostate cancer is associated with reduced fraction of CD8 effector cells and increased TIGIT"

**Figure S1**

A

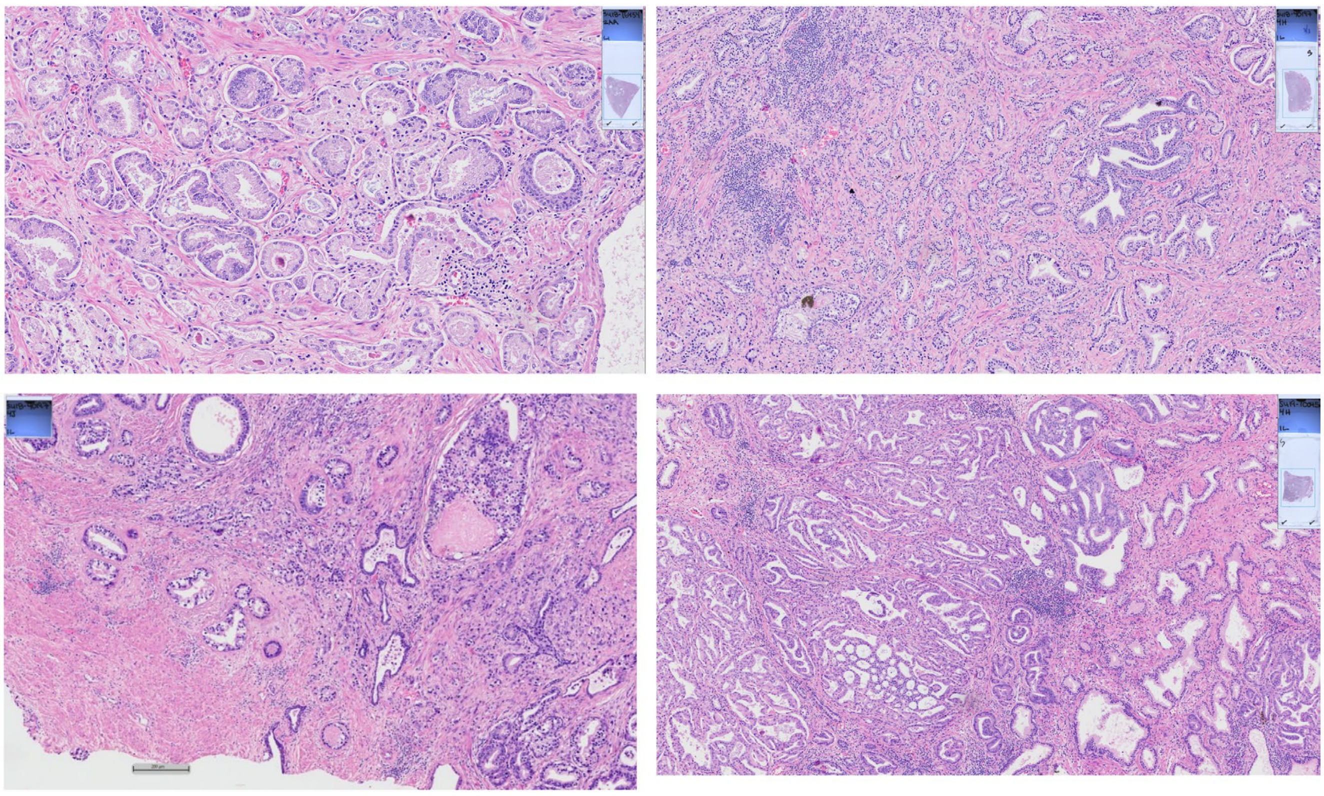

B

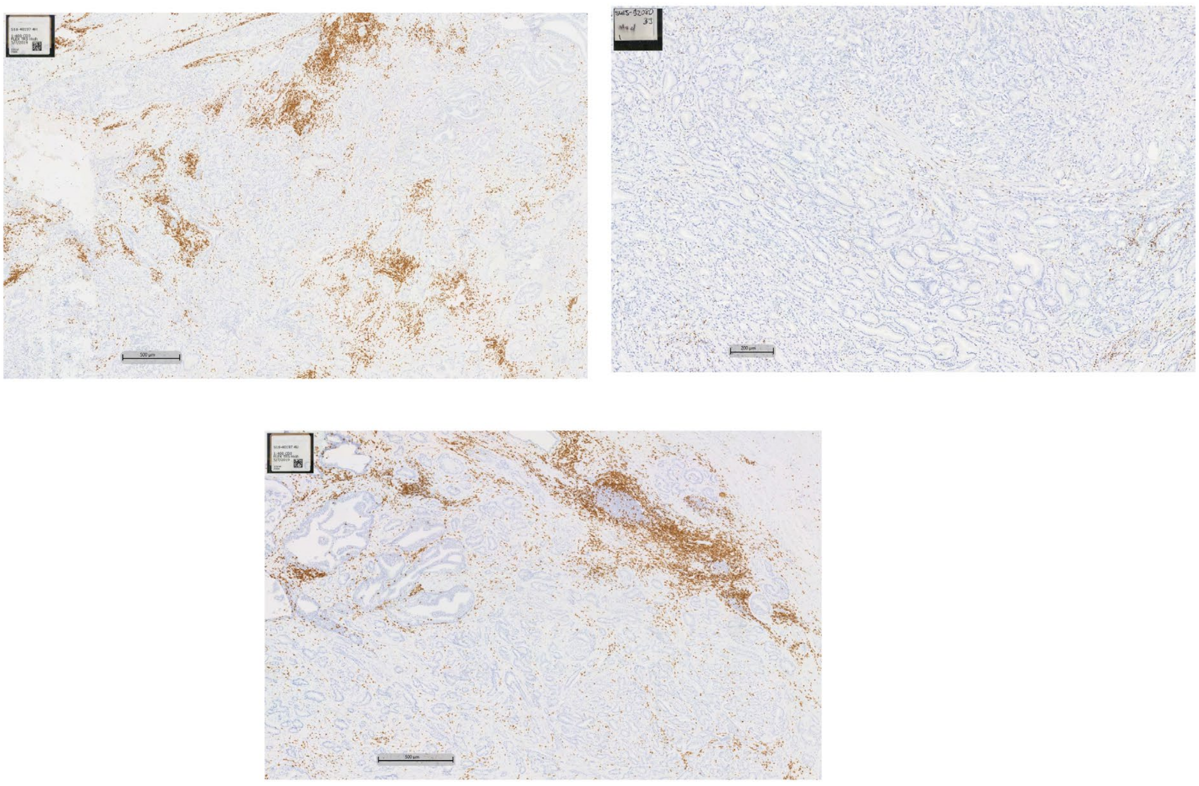

**Figure S1. Histologic examples of low and high T-cell infiltration in untreated primary prostate cancer.** (A) Representative hematoxylin-and-eosin-stained sections. (B) Representative CD3 immunohistochemistry illustrating the corresponding range of T-cell infiltration.

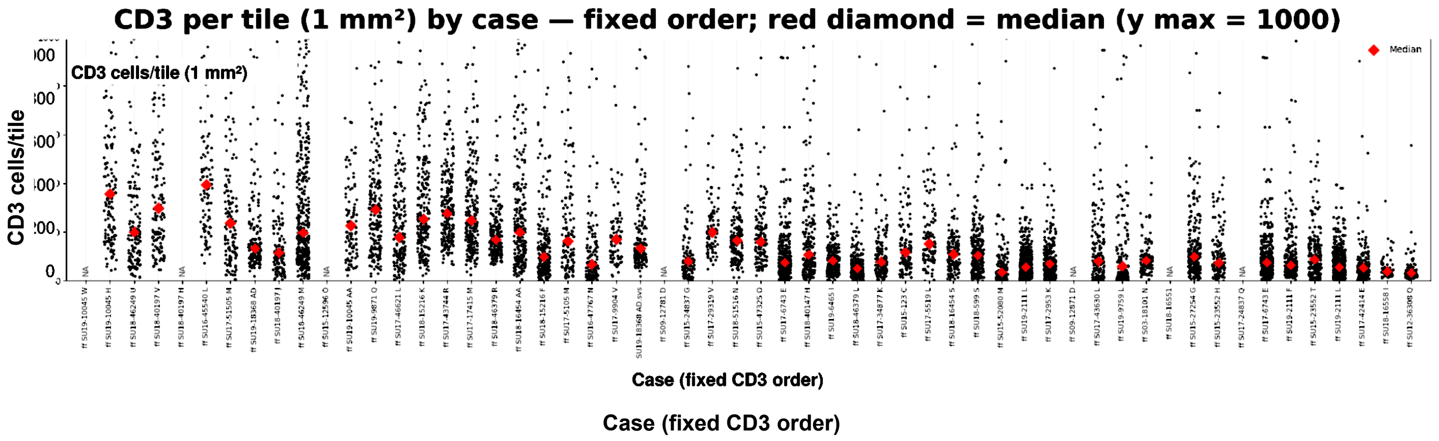


**Figure S2. Expanded tile-level distribution of CD3⁺ T-cell density.** CD3⁺ cell density is shown for every evaluable 1-mm² tile within each case. Each point represents one tile, red diamonds indicate the case median, and cases are ordered from highest to lowest whole-slide CD3⁺ density. The y-axis is truncated at 1,000 cells/mm² for visualization; higher-density tiles are retained at the upper plotting limit. Cases with unavailable measurements are indicated as not available.

### Figure S3

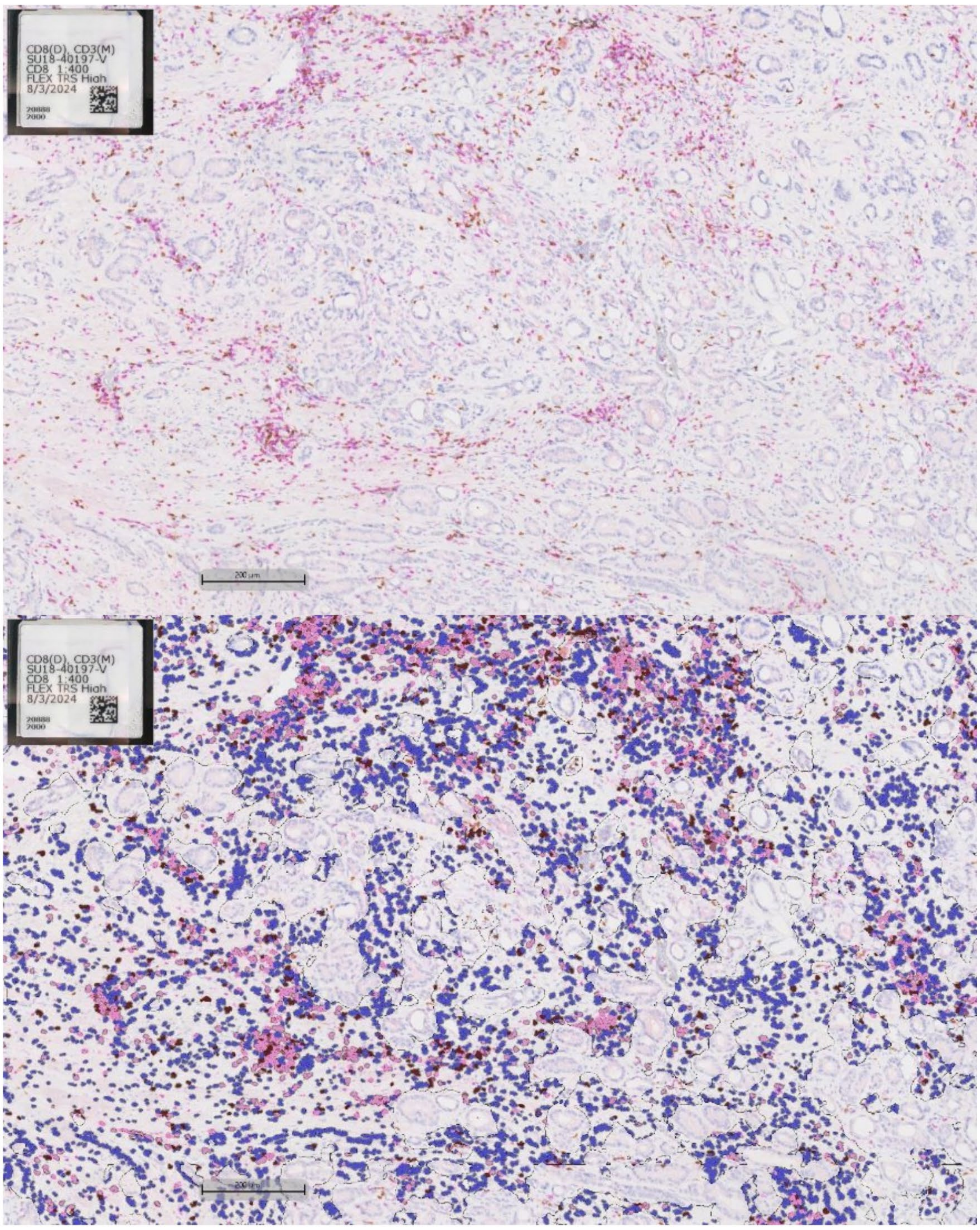


**Figure S3. Representative dual CD3/CD8 immunohistochemistry.** Representative low- and high-density T-cell infiltrates are shown using dual chromogenic staining for CD3 and CD8.

Figure S4

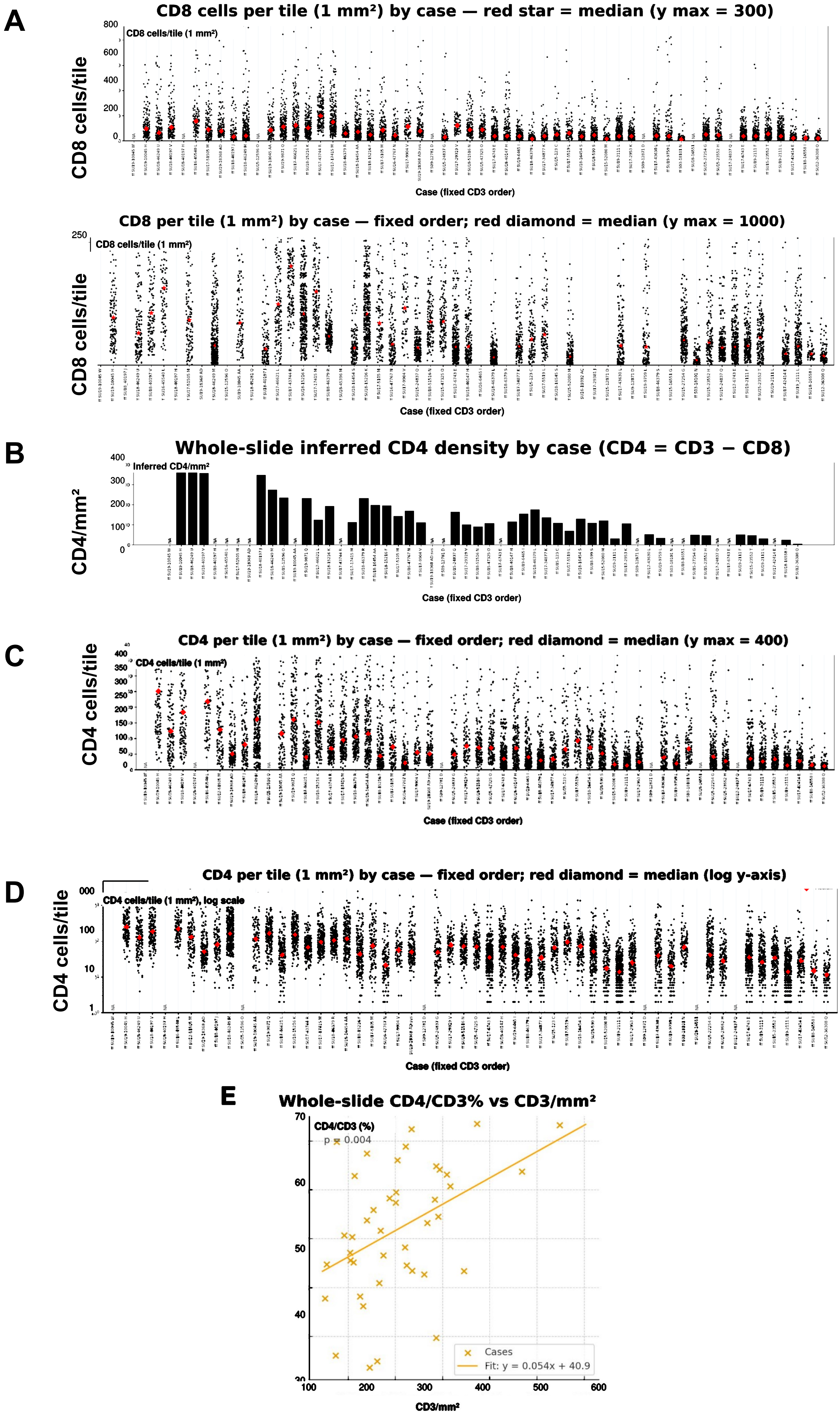


**Figure S4. Spatial distribution of CD8⁺ and inferred CD4⁺ T cells.** (A) Tile-level CD8⁺ cell density across cases, with each point representing one 1-mm² tile and red diamonds indicating case medians; cases are ordered by whole-slide CD3⁺ density. (B) Inferred whole-slide CD4⁺ cell density by case. CD4⁺ abundance was calculated as CD3⁺ minus CD8⁺ cells. (C, D) Tile-level inferred CD4⁺ cell density displayed across cases; points represent individual 1-mm² tiles and red diamonds indicate case medians. (E) Positive relationship between the inferred whole-slide CD4/CD3 ratio and whole-slide CD3⁺ density (Pearson $r = 0.43$, $p = 0.004$); the line shows the linear fit.

### Figure S5

**A** PD1 IHC

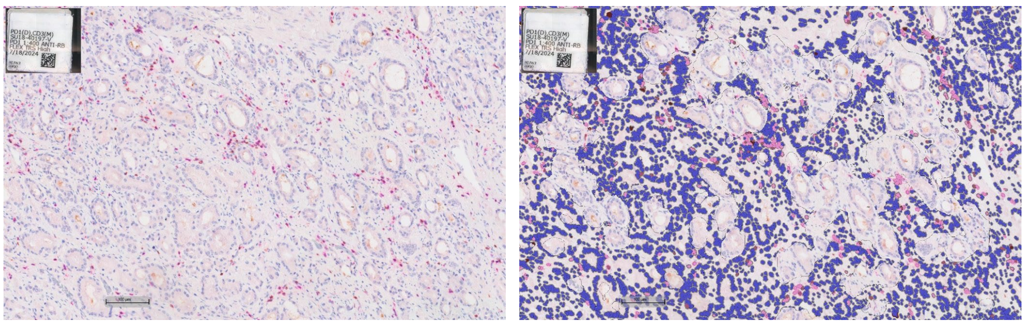

**B**

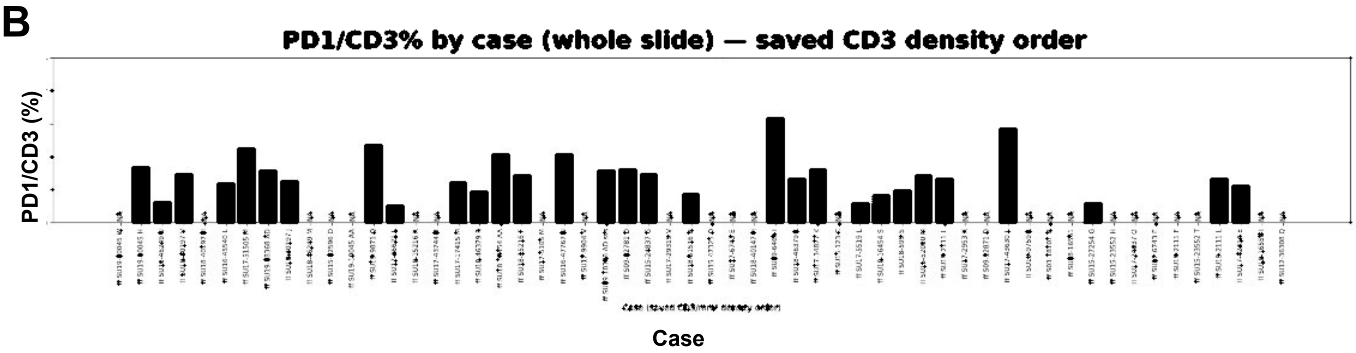


**C**

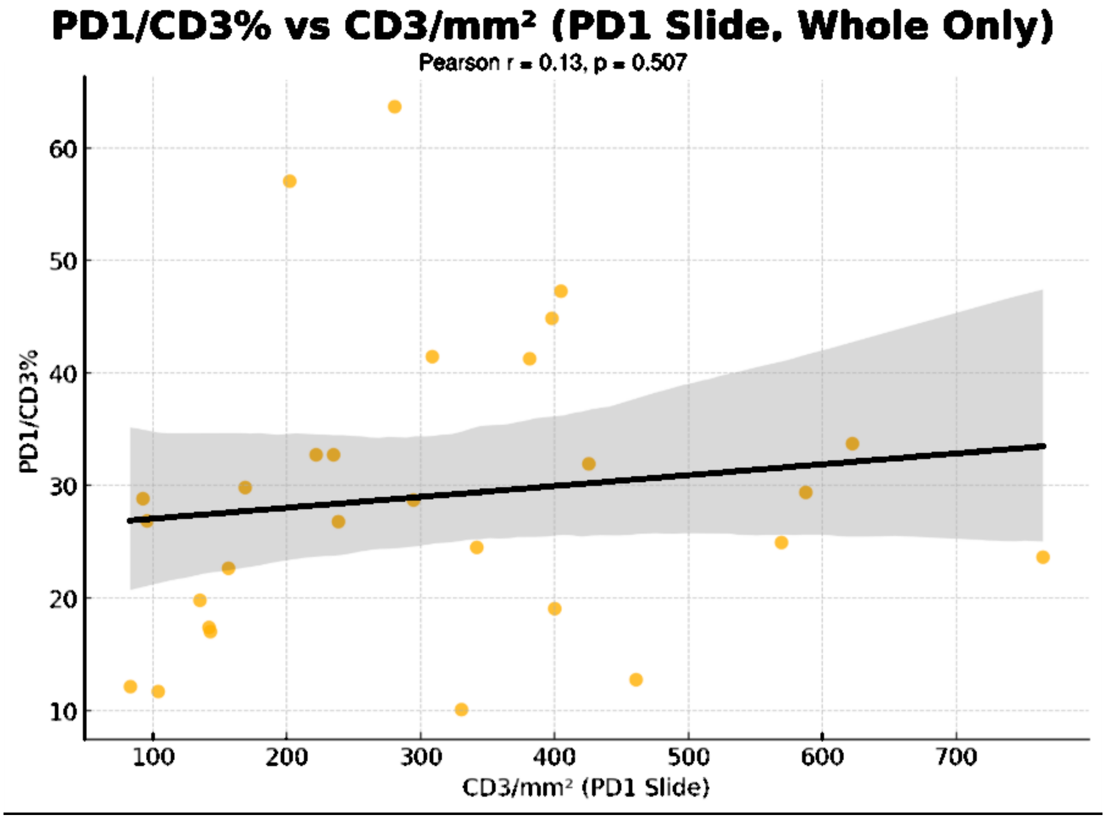


**Figure S5. PD1 expression is not associated with overall T-cell density.** (A) Representative PD1 immunohistochemistry. (B) Whole-slide PD1/CD3 ratio by case, with cases ordered by whole-slide $CD3^+$ density; unavailable measurements are indicated as not available. (C) Association between PD1/CD3 and whole-slide $CD3^+$ density (Pearson $r = 0.13$, $p = 0.507$); the line shows the linear fit and shading shows the 95% confidence interval.

### Figure S6

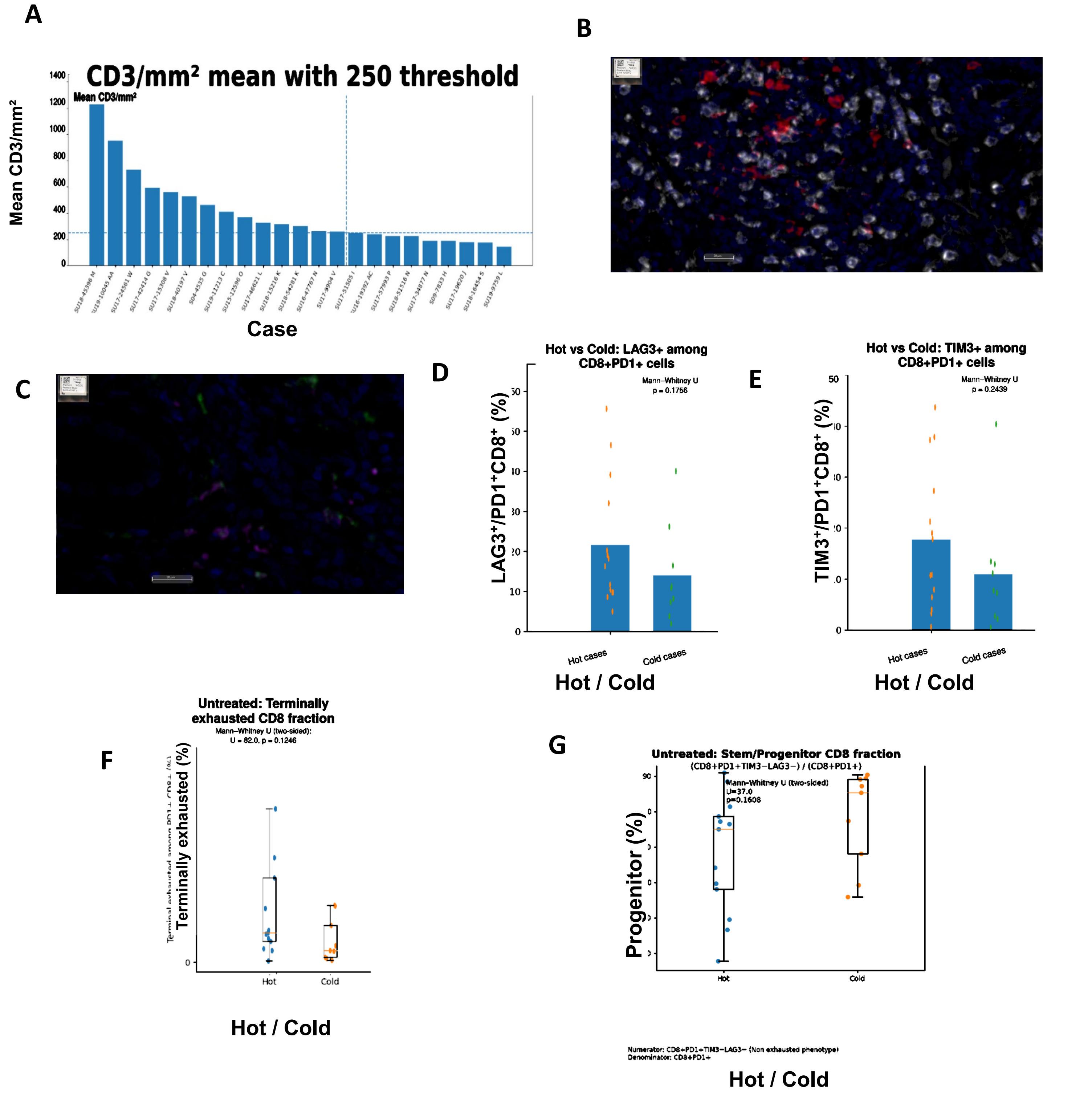


**Figure S6. Additional immune phenotypes in Hot and Cold tumors.** (A) Whole-slide CD3⁺ density and the prespecified 250-cells/mm² threshold used to classify tumors as Hot or Cold. Representative multiplex immunofluorescence images show (B) CD8 (displayed in white; Opal 480) with PD1 (displayed in red; Opal 690) and (C) CD163 (displayed in purple; Opal 780) with C1Q (displayed in green; Opal 520). Within $PD1^+CD8^+$ T cells, panels show the percentages expressing (D) LAG3, (E) TIM3, (F) both TIM3 and LAG3, and (G) neither TIM3 nor LAG3. Box plots show the median and interquartile range. Hot-versus-Cold comparisons used two-sided Mann–Whitney U tests; none of the displayed comparisons reached statistical significance.

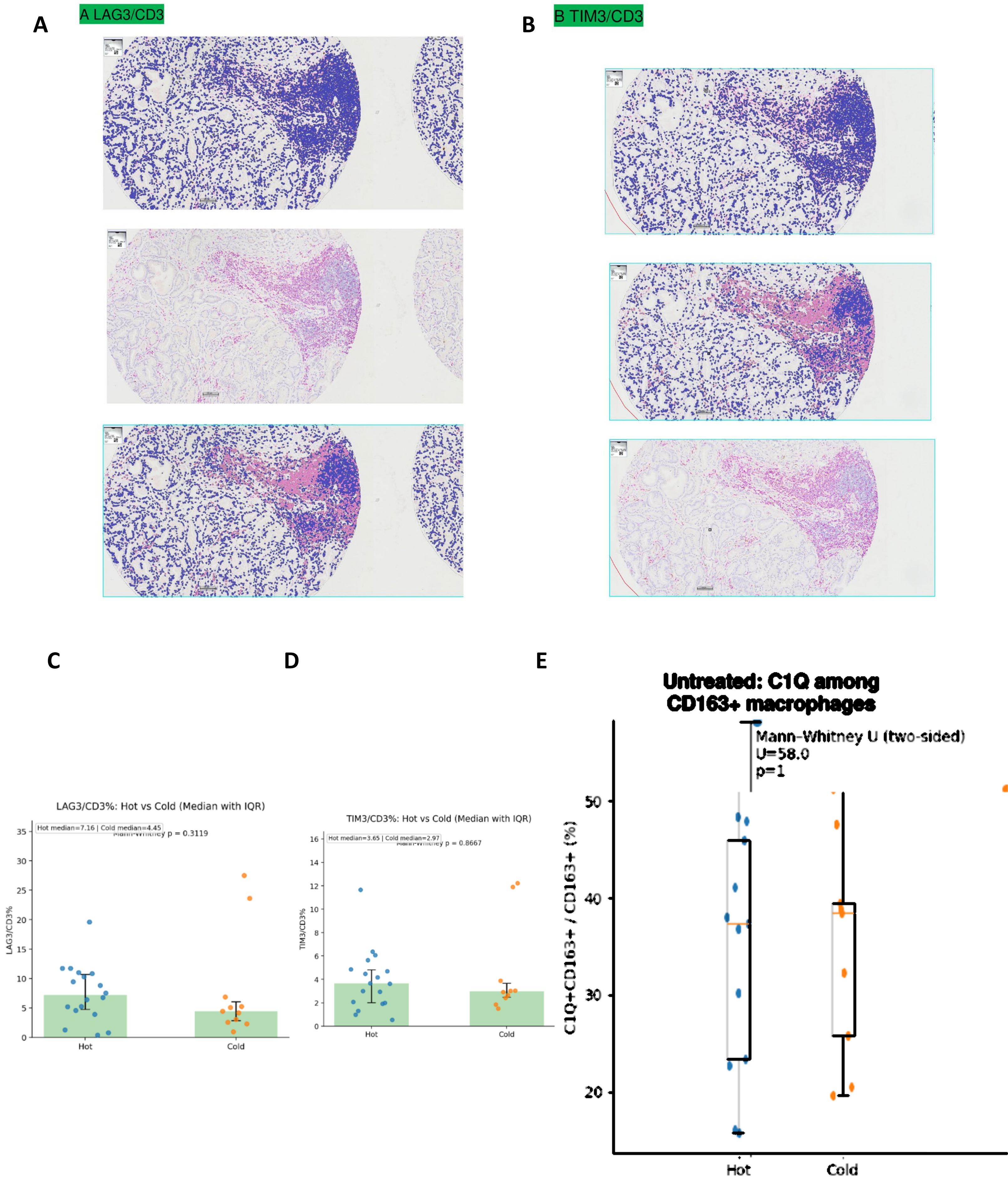


**Figure S7. LAG3, TIM3, and macrophage C1Q features in Hot and Cold tumors.** (A) Representative dual chromogenic LAG3/CD3 immunohistochemistry. (B) Representative dual chromogenic TIM3/CD3 immunohistochemistry. (C) LAG3/CD3 in Hot and Cold tumors. (D) TIM3/CD3 in Hot and Cold tumors. (E) C1Q$^{+}$CD163$^{+}$ cells as a percentage of CD163$^{+}$ macrophages in Hot and Cold tumors. For panels C and D, points represent individual tumors, bars show medians with interquartile-range error bars, and two-sided Mann–Whitney U-test results are displayed. Panel E is displayed as indicated.

Figure S8

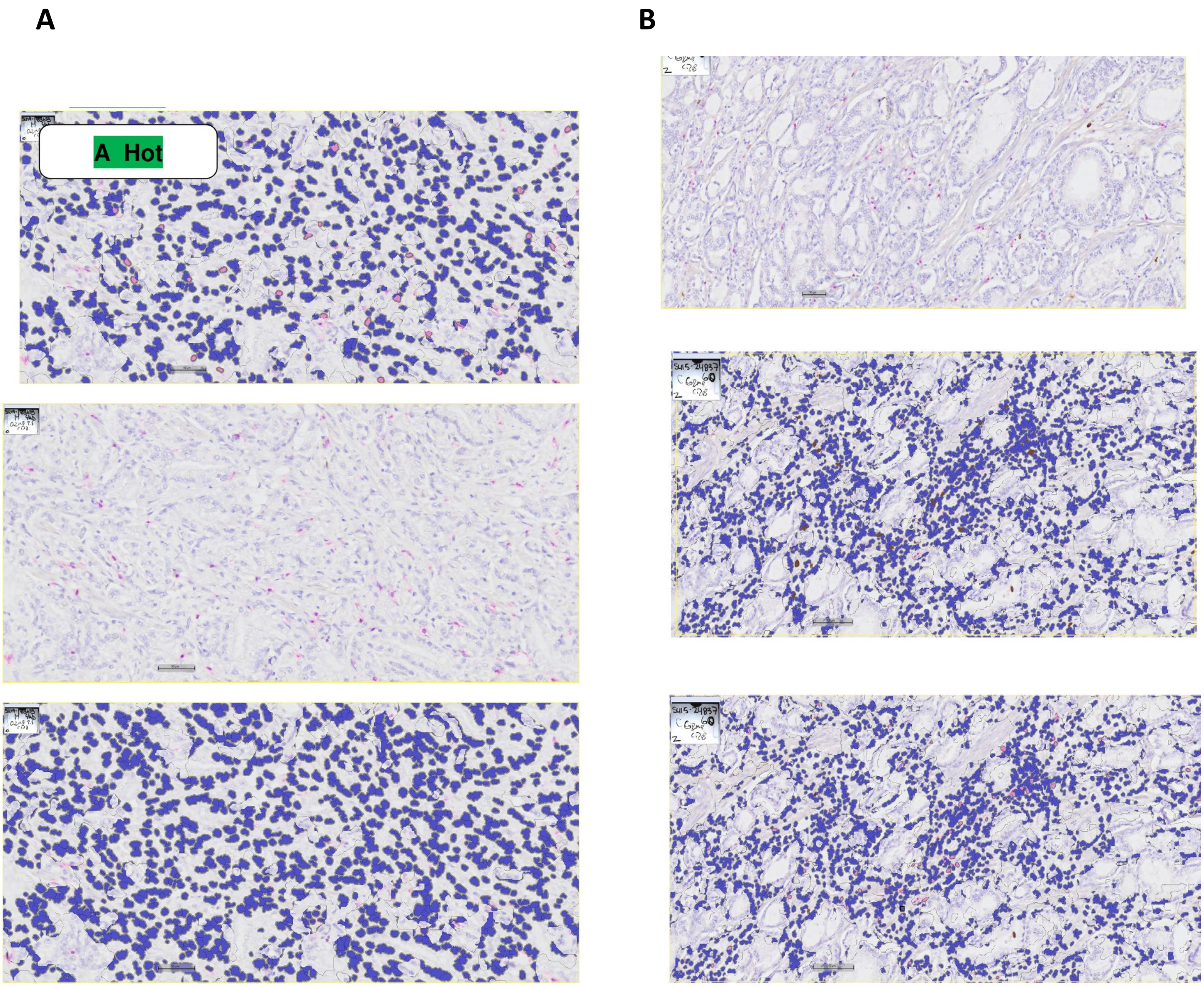


**Figure S8. Representative dual GZMB/CD8 immunohistochemistry in Hot and Cold tumors.** (A) Hot tumor and (B) Cold tumor, classified according to whole-slide $CD3^{+}$ T-cell density. The representative cold case shows a higher GZMB/CD8 ratio than the representative hot case. GZMB was visualized with DAB and CD8 with Magenta chromogen.

### Figure S9

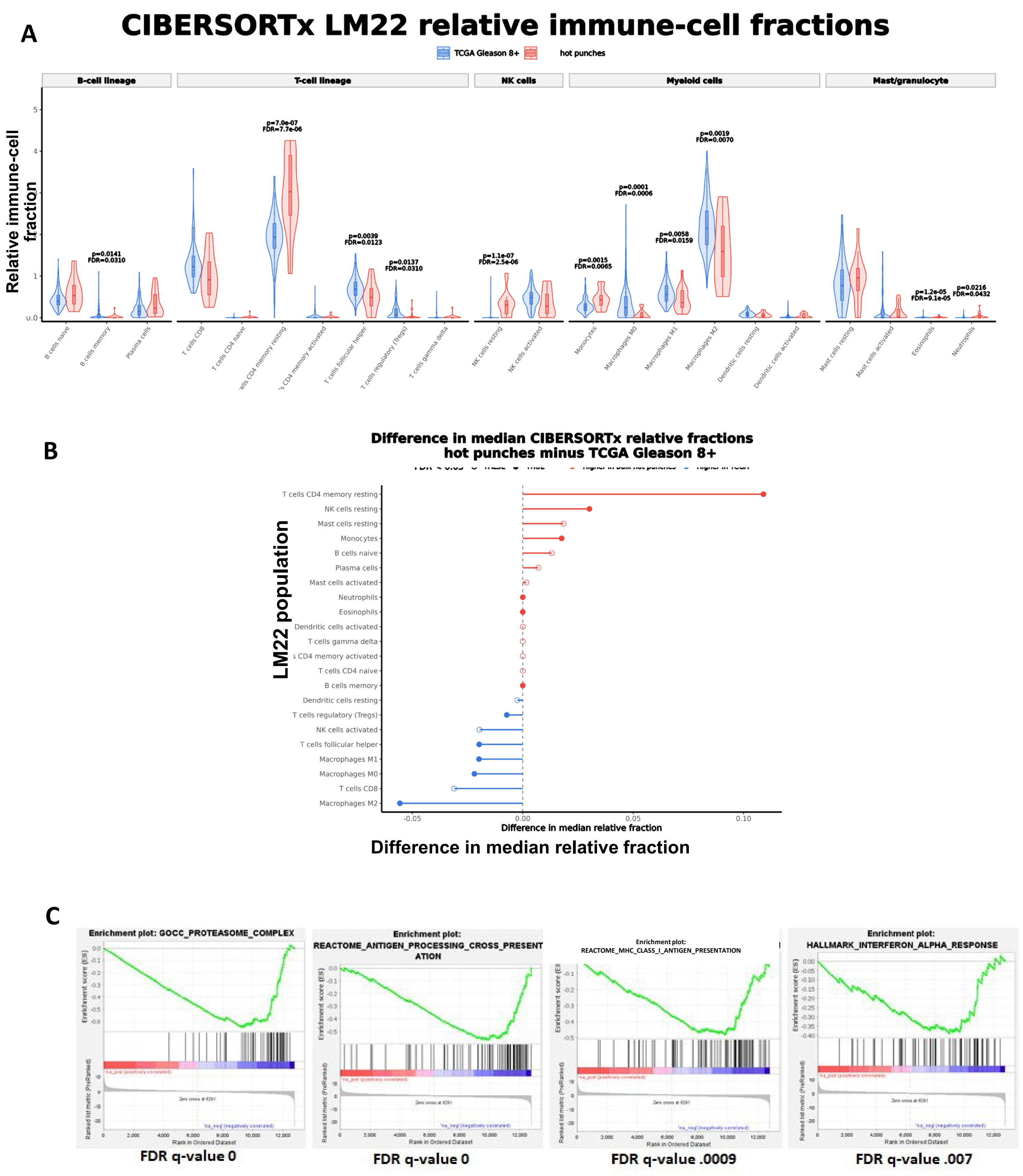


**Figure S9. CIBERSORTx immune-cell composition and gene-set enrichment in Hot punches.** CIBERSORTx LM22 relative immune-cell fractions were compared between 19 evaluable Hot punches and 193 TCGA primary prostate cancers with Gleason score ≥8. (A) Relative fractions for all 22 LM22 immune populations. (B) Difference in median relative fraction between cohorts for each LM22 population; positive values indicate higher relative representation in the Hot punches and negative values indicate higher representation in TCGA. Comparisons used two-sided nonparametric tests with Benjamini–Hochberg correction. Because LM22 fractions sum to 100% within each sample, results reflect relative compositional redistribution rather than absolute cell abundance. (C) Gene-set enrichment analysis for GOCC_PROTEASOME_COMPLEX, REACTOME_ANTIGEN_PROCESSING_CROSS_PRESENTATION, REACTOME_MHC_CLASS_I_ANTIGEN_PRESENTATION, and HALLMARK_INTERFERON_ALPHA_RESPONSE; the displayed FDR q values are 0, 0, 0.0009, and 0.007, respectively.

### Figure S10

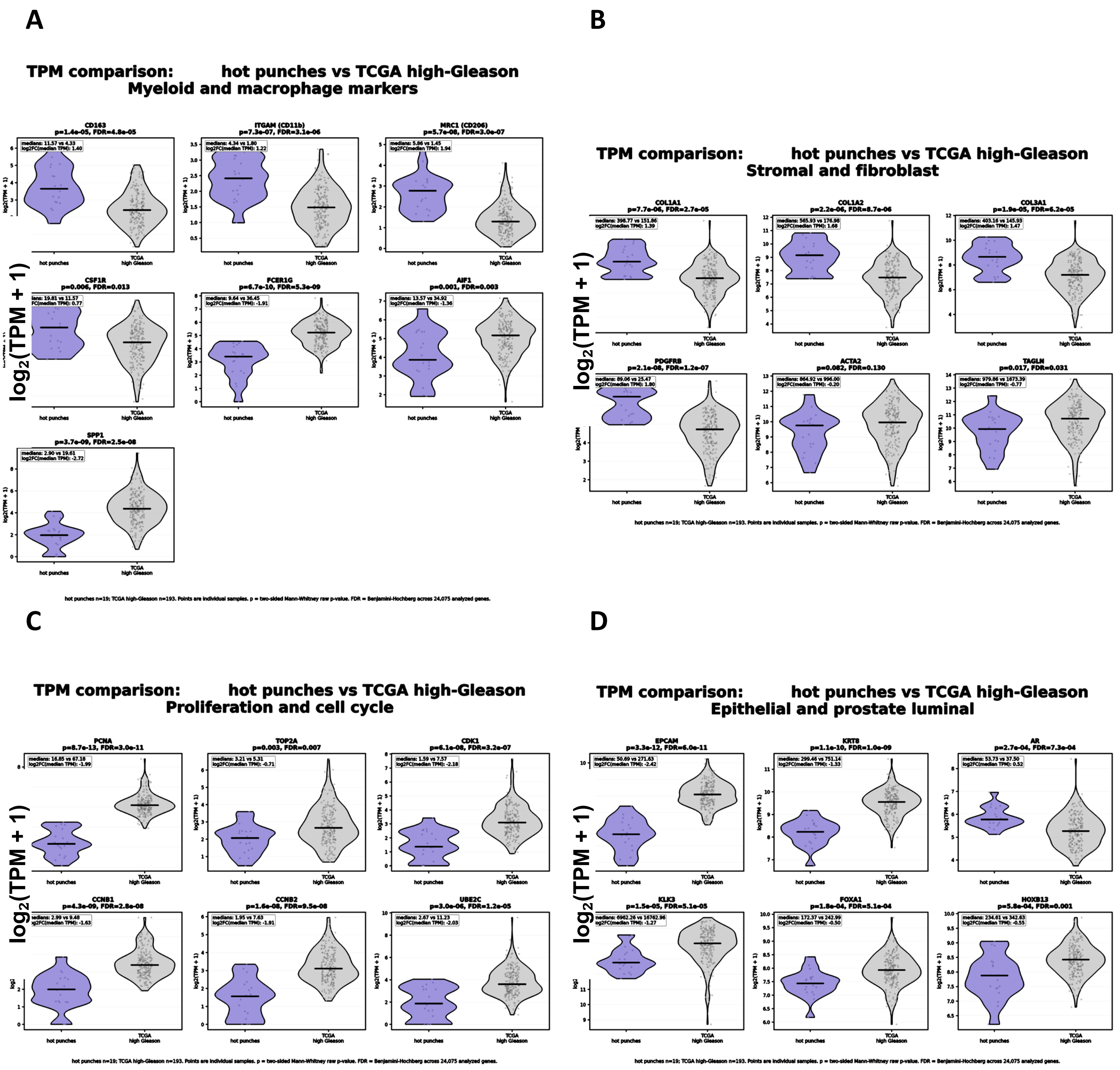


**Figure S10. Additional bulk RNA-sequencing comparisons of Hot punches with high-Gleason TCGA tumors.** Transcript abundance in 19 evaluable Hot punches was compared with 193 TCGA primary prostate cancers with Gleason score ≥8 after matched RSEM processing. Violin plots show transcripts per million for (A) myeloid and macrophage markers, (B) stromal and fibroblast genes, (C) proliferation and cell-cycle genes, and (D) epithelial and prostate-luminal genes. Points represent individual samples and horizontal lines indicate distribution summaries. Two-sided between-cohort tests were corrected for multiple comparisons as specified in the Methods.

### Figure S11

A

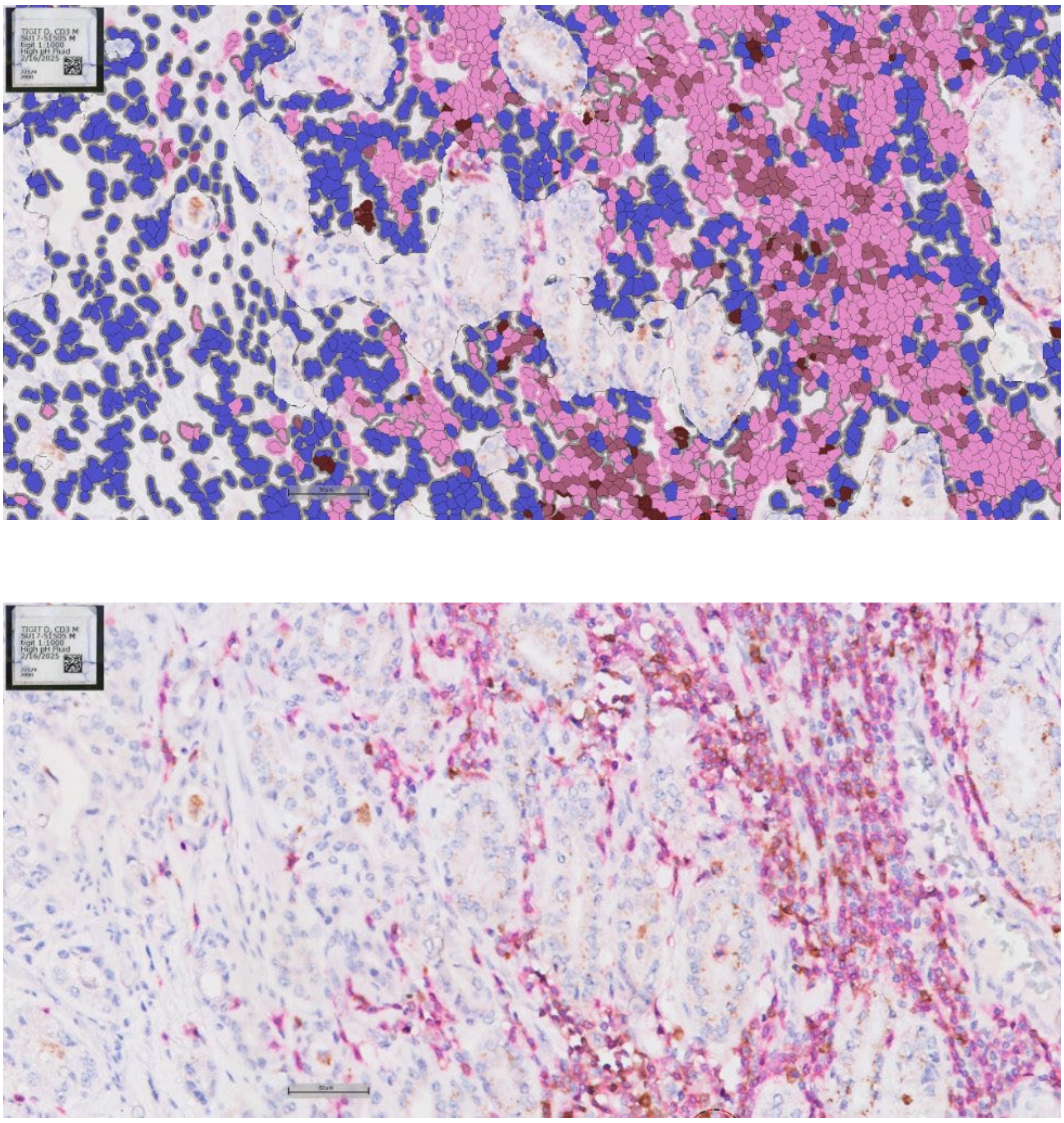

B

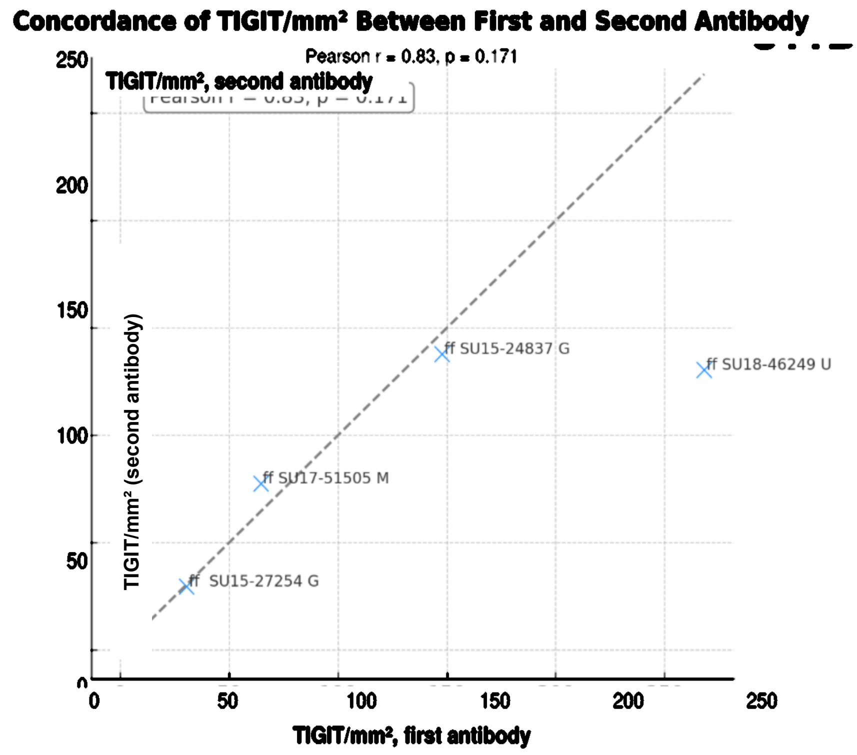


**Figure S11. TIGIT immunohistochemistry and independent-antibody validation.** (A) Representative TIGIT immunohistochemistry. (B) Concordance of TIGIT⁺ cell density measured using the original anti-TIGIT antibody (BLR047F, Fortis Life Sciences) and a second anti-TIGIT antibody (ab243903, Abcam) in the available validation cases. The dashed line indicates identity (Pearson r = 0.83, p = 0.171).

### Figure S12

**A**

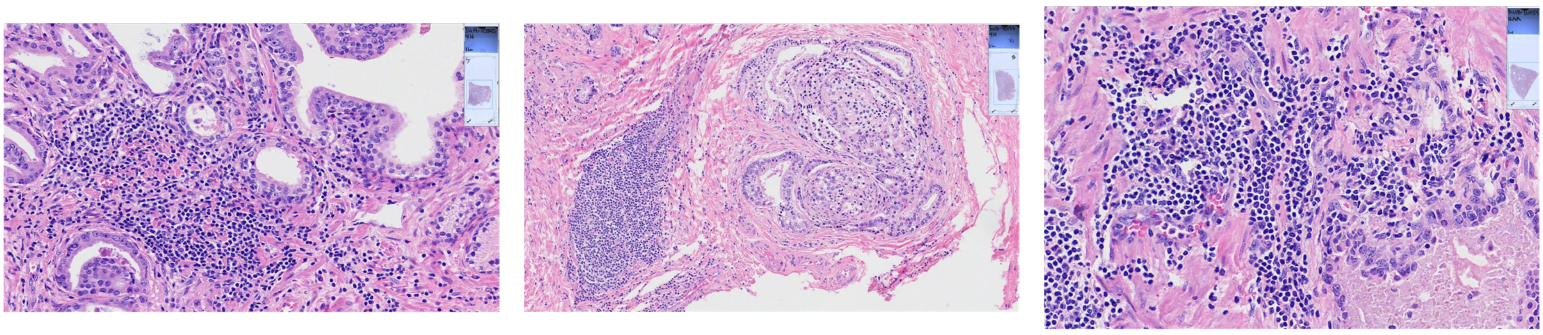

**B**

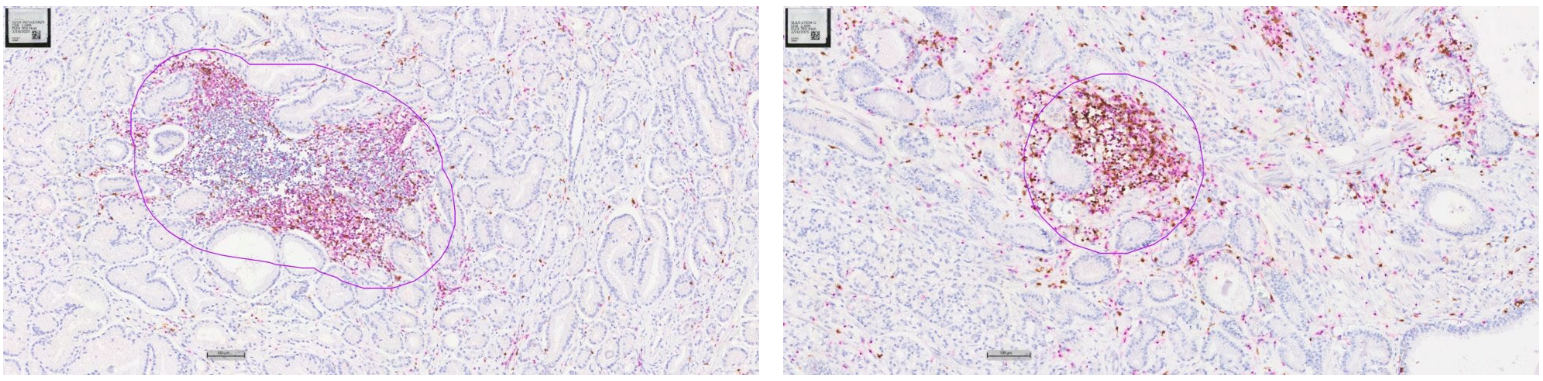

**Figure S12. Histologic features of lymphoid aggregates.** Lymphoid aggregates were defined as regions containing >3000 $CD3^{+}$ cells/$mm^{2}$. (A) Representative hematoxylin-and-eosin-stained lymphoid aggregates. (B) Representative dual chromogenic CD3/CD8 immunohistochemistry showing predominantly T-cell-rich aggregates, with CD3 visualized using DAB and CD8 using Magenta chromogen. These aggregates were not established as mature tertiary lymphoid structures.
